# An *in silico* method to assess antibody fragment polyreactivity

**DOI:** 10.1101/2022.01.12.476085

**Authors:** Edward P. Harvey, Jung-Eun Shin, Meredith A. Skiba, Genevieve R. Nemeth, Joseph D. Hurley, Alon Wellner, Ada Y. Shaw, Victor G. Miranda, Joseph K. Min, Chang C. Liu, Debora S. Marks, Andrew C. Kruse

## Abstract

Antibodies are essential biological research tools and important therapeutic agents, but some exhibit non-specific binding to off-target proteins and other biomolecules. Such polyreactive antibodies compromise screening pipelines, lead to incorrect and irreproducible experimental results, and are generally intractable for clinical development. We designed a set of experiments using a diverse naïve synthetic camelid antibody fragment (‘nanobody’) library to enable machine learning models to accurately assess polyreactivity from protein sequence (AUC > 0.8). Moreover, our models provide quantitative scoring metrics that predict the effect of amino acid substitutions on polyreactivity. We experimentally tested our model’s performance on three independent nanobody scaffolds, where over 90% of predicted substitutions successfully reduced polyreactivity. Importantly, the model allowed us to diminish the polyreactivity of an angiotensin II type I receptor antagonist nanobody, without compromising its pharmacological properties. We provide a companion web-server that offers a straightforward means of predicting polyreactivity and polyreactivity-reducing mutations for any given nanobody sequence.

## INTRODUCTION

Due to their specificity and affinity, antibodies are an indispensable class of biomedical research tools as well as important therapeutics for the treatment of cancer, autoimmune, and infectious diseases. Current antibody discovery methods prioritize the generation of antibodies and antibody fragments with high target specificity. However, some antibodies that strongly bind one target interact with additional antigens with low-affinity. In clinical development, these non-specific or polyreactive antibodies show poor pharmacokinetics or other liabilities that limit clinical use^1-3^. Additionally, polyreactive antibodies encountered in the basic research setting cause misinterpretation of results, low reproducibility in routine experiments, and wasted time and money^4^. Thus, there have been several calls to standardize the quality and specificity of antibodies used in research settings similar to those in the clinic^5,6^.

Developing and improving methods to detect and quantify polyreactivity are essential for enhancing the quality of antibodies in both clinical development and basic research settings. Many experimental methods that evaluate polyreactivity^7-14^ are low-throughput and require experimental screening with purified antibody. The degree of polyreactivity is highly method and reagent-dependent and is typically measured after antigen selection, making it difficult to prioritize the most promising clones. Understanding sequence features of polyreactive antibodies could provide an efficient avenue to quantitatively assess antibody polyreactivity without experimental effort. Previous computational methods^15-22^ have revealed features of polyreactivity antibodies, such as J- and V-chain usage^17^, high isoelectric points in the complementarity determining regions (CDRs)^16,18-25^, longer CDR3s^16,23^, enrichment of arginine, glycine, valine, and tryptophan containing motifs^18^, and glutamine residues^23^. Despite these extensive analyses the relative importance of many characteristics is disputed^21^ and prediction software cannot quantitate polyreactivity^17^.

For broad utility, a computational method should accurately predict the *degree* of polyreactivity and compute candidate rescue mutations from the input of a user sequence alone. To achieve this goal, we designed experiments to learn features of high and low polyreactivity clones from a naïve synthetic yeast display library of heavy-chain only camelid antibody fragments (nanobodies)^26,27^ through computational methods. Synthetic nanobodies provide an ideal reductionist system to probe polyreactivity in the context of a fixed framework without the influence of heavy and light chain pairing effects. These methods result in generalizable software that quantifies nanobody polyreactivity based on sequence alone and most importantly designs specific mutations to decrease polyreactivity.

We successfully applied our software to three polyreactive nanobodies, including AT118i4h32, a nanobody antagonist of the angiotensin II type I receptor (AT1R)^28^, where we reduced polyreactivity without compromising binding affinity or target-specific pharmacology. This sequence-based approach may be a generally useful tool for prioritizing nanobody clones identified in selection experiments and improving nanobodies targeting diverse antigens. While nanobodies are gaining popularity as next generation biotherapeutics^29^ that target antigen surfaces and tissue types not accessible to conventional antibodies, the approaches developed here are in principle fully applicable to conventional antibodies as well.

## RESULTS

### Enriching naïve library for polyreactive clones

Unlike previous analyses of antibody polyreactivity which relied on clinical candidates ^23-25^, clones enriched for antigen binding^17^, or primarily focused on the contribution of V_H_ CDR3 antibody polyreactivity^18,21^, we designed experiments to assess polyreactivity of clones from a naïve synthetic yeast display library through binding to detergent-solubilized *Spodoptera frugiperda* (Sf9) insect cell membranes (Figure 1)^14^. This mixed protein polyspecificity reagent (PSR) is compatible with sorting large pools of antigen naïve clones, allowing us to determine global contributions to polyreactivity in an unbiased manner. The yeast display library contains >,2×10^9^ unique nanobody clones that mimic a naïve llama immune repertoire in CDR sequence composition and CDR3 length and possesses moderate diversity in the CDR1 and CDR2 regions and extensive diversity in the CDR3 region. We used Magnetic-Activated Cell Sorting (MACS) to both enrich for polyreactive clones and deplete non-expressing clones from the library. Following MACS, distinct populations of clones with high and low polyreactivity were isolated by Fluorescence-Activated Cell Sorting (FACS) (Supplementary Figure 1A-B).

**Figure 1.**
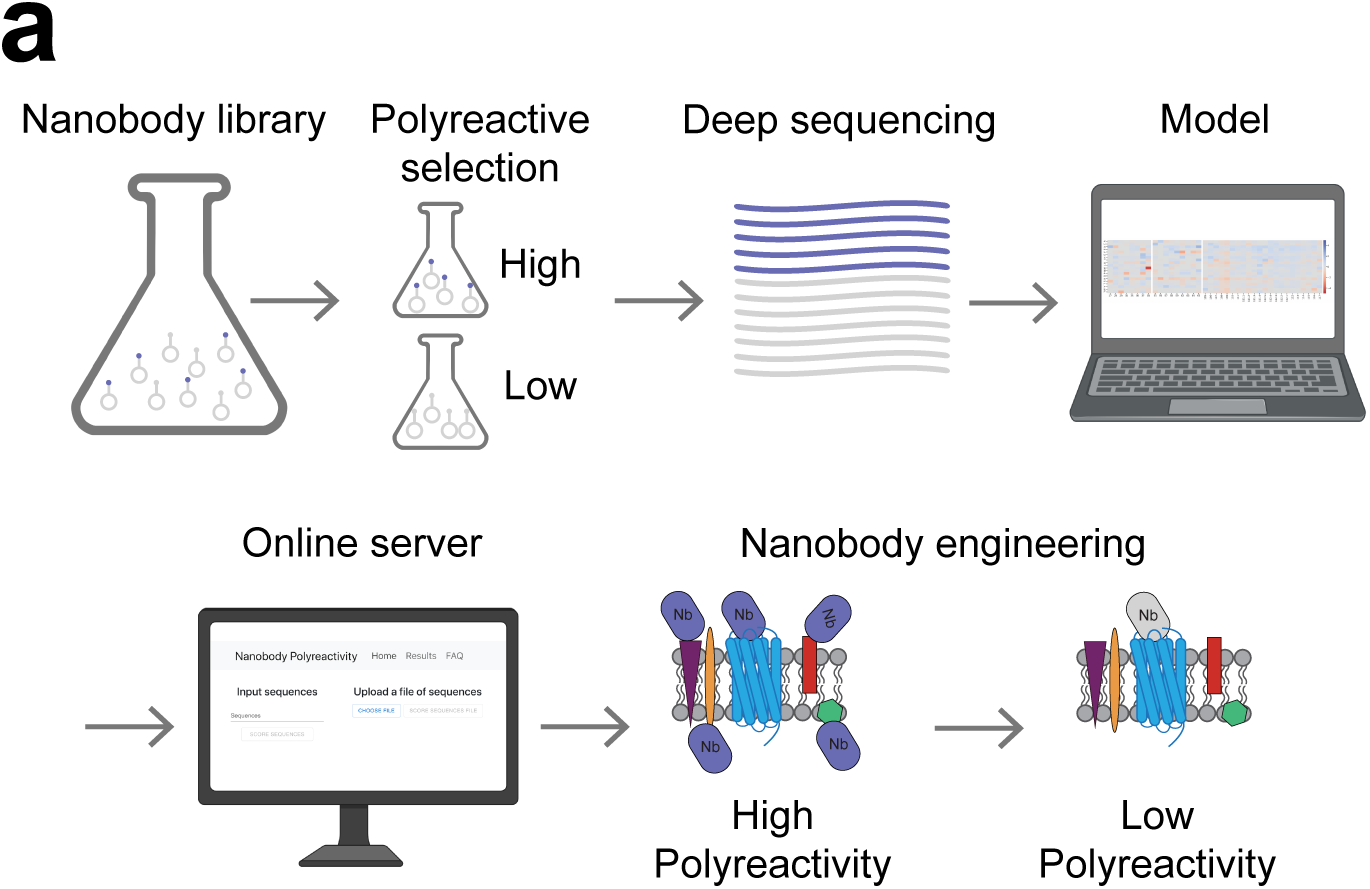
Development of computational tool to assess and mitigate polyreactivity. Starting from a large, naïve synthetic nanobody library, pools of nanobodies with low and high polyreactivity were isolated. Machine learning models were trained on deep sequencing data from these pools to learn sequence features of low and high polyreactive nanobodies. These algorithms were incorporated into software that quantitatively predicts polyreactivity levels and recommends substitutions that reduce it.

PSR reagent has not been used to assess nanobody polyreactivity, but is well validated against other measures of polyreactivity for conventional antibodies^2,14,15^. To validate PSR performance on nanobodies, we recombinantly expressed six nanobodies with varying levels of polyreactivity from our FACS sorted pools and assessed polyreactivity by conventional ELISA assays against lysozyme, double stranded DNA (dsDNA), single stranded DNA (ssDNA), insulin, lipopolysaccharide (LPS), and bare plastic (Figure 2, Supplementary Figure 2A-F). ELISA polyreactivity assays performed using different reagents correlated well with one another (r^2^ values between 0.789 and 0.986, p < 0.05) with the exception of lysozyme (r^2^ values between -0.109 and 0.045, p-values between 0.8127 and 0.9230), which did not correlate with the other reagents. Furthermore, direct ELISA assays strongly correlated with insect cell PSR (r^2^ values between 0.7849 and 0.9268) except for lysozyme which exhibited a very weak correlation (r^2^ = -0.1876). The correlations between insulin, LPS, and ssDNA direct ELISA assays to insect cell PSR staining were highly significant (p < 0.05), while bare plastic and dsDNA direct ELISA assays were modestly significant (p < 0.10). Lysozyme direct ELISA assays did not significantly correlate with insect cell PSR staining (p = 0.7219). We also observed that polyreactive clones had increased retention times in conventional size exclusion chromatography albeit not with statistical significance (r^2^ = 0.7836, p = 0.1168), suggesting that nanobody polyreactivity may be detected during routine protein purification (Supplementary Figure 2G). Overall, the ELISA experiments support that the pools of nanobodies selected by PSR staining possess high and low levels of polyreactivity. Armed with this validation, we deep-sequenced the two FACS sorted pools and obtained 65,147 unique low polyreactivity sequences and 69,155 unique highly polyreactive sequences that contained 51,308 and 59,623 distinct CDR regions.

**Figure 2.**
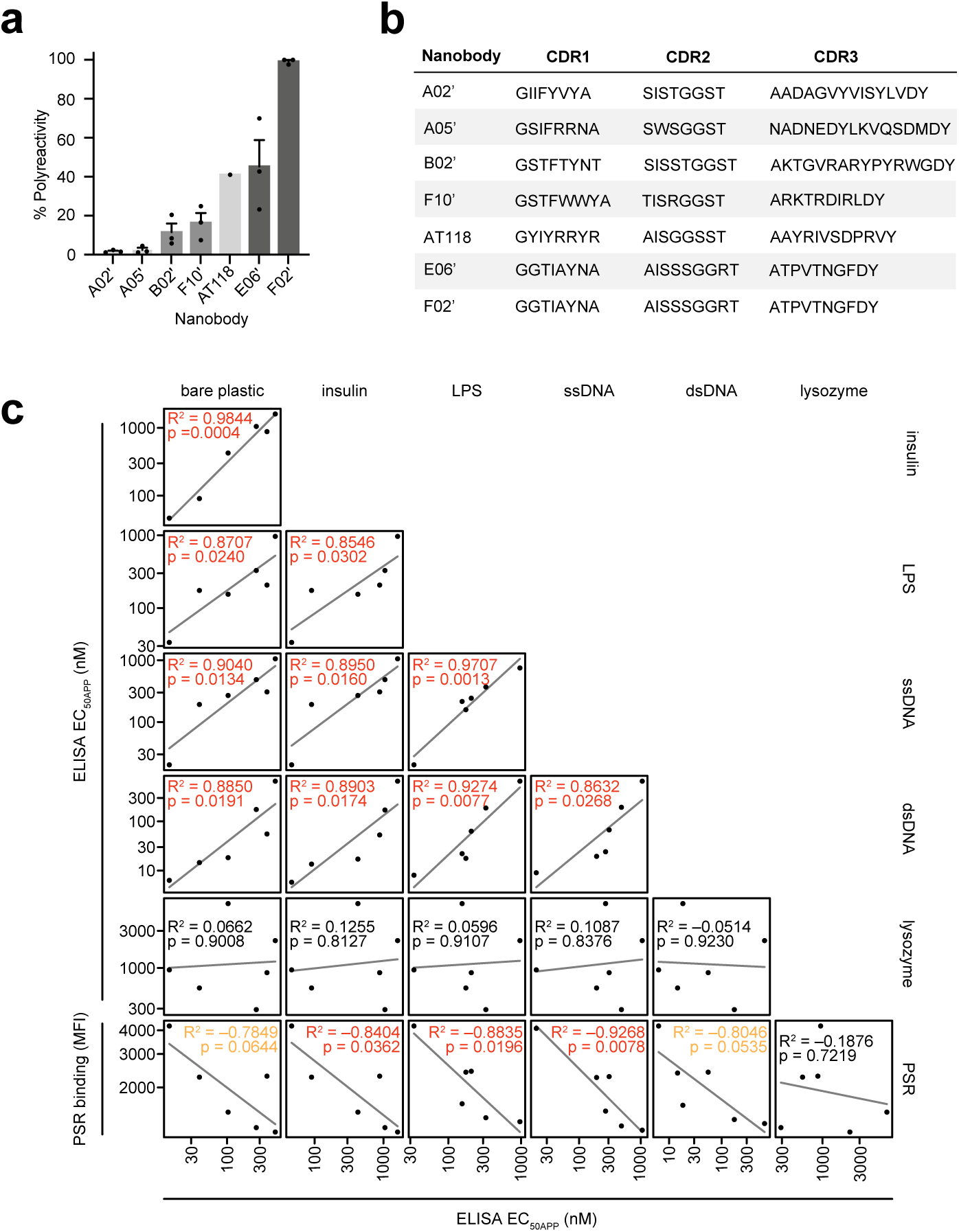
Correlations between direct ELISA assays and insect cell polyspecificity reagent (PSR) staining. **a**, *Spodoptera frugiperda* (Sf9) insect cell PSR staining of single nanobodies isolated from FACS sorts. Data are mean +/- SEM of three independent biological experiments performed in technical triplicate. Polyreactivity levels are normalized with respect to the highest value. **b**, CDR sequences of isolated nanobodies. **c**, Direct ELISA assays measured the apparent EC_50_ (EC_50APP_) of five index panel members and nanobody AT118 to the specified reagents. ELISA data are representative of two independent experiments, each performed in technical triplicates.

### Development of computational method

We developed computational models trained on the sequences from the FACS-sorted pools to classify nanobodies as possessing high or low polyreactivity. We constructed a suite of supervised, discriminative models that can separate high and low polyreactivity sequences (Figure 3A-B). These models include a logistic regression model of a one-hot embedding of the CDR sequences, a logistic regression model of a k-mer embedding (k=3) of the CDR sequences, a convolutional neural network (CNN), and a recurrent neural network (RNN). The one-hot logistic regression model learns weights for each amino acid type at each position in the CDR sequences that are most predictive of polyreactivity; the k-mer logistic regression learns weights for each motif (lengths 1, 2, and 3) that are most predictive of polyreactivity, irrespective of where they occur within a given CDR sequence. Convolutional neural networks use convolutional filters to learn spatial information (e.g., an amino acid and its neighboring residues) and are often used in image classification. Recurrent neural networks capture sequential information (e.g. the probability of a residue given the previous residues) and are frequently used in text and audio analysis. For the one-hot logistic regression and for the CNN, we align the CDR sequences using the IMGT numbering scheme with ANARCI^30^. The k-mer logistic regression and the RNN methods do not require aligned CDR sequences. In order to test the generalizability of our models, we clustered the nanobody sequences using k-means clustering to generate five clusters of sequences, which we used to build train and test splits. These splits and careful selection allowed us to avoid over-optimistic prediction accuracies that result from the tests sets overlapping or close to the training sets^31^. Specifically, we ensured that all sequences in the test sets were more than 10 edit-distance (Levenshtein distance) and possessed only ∼75% sequence similarity in the CDR sequences from each other (Figure 3A).

**Figure 3.**
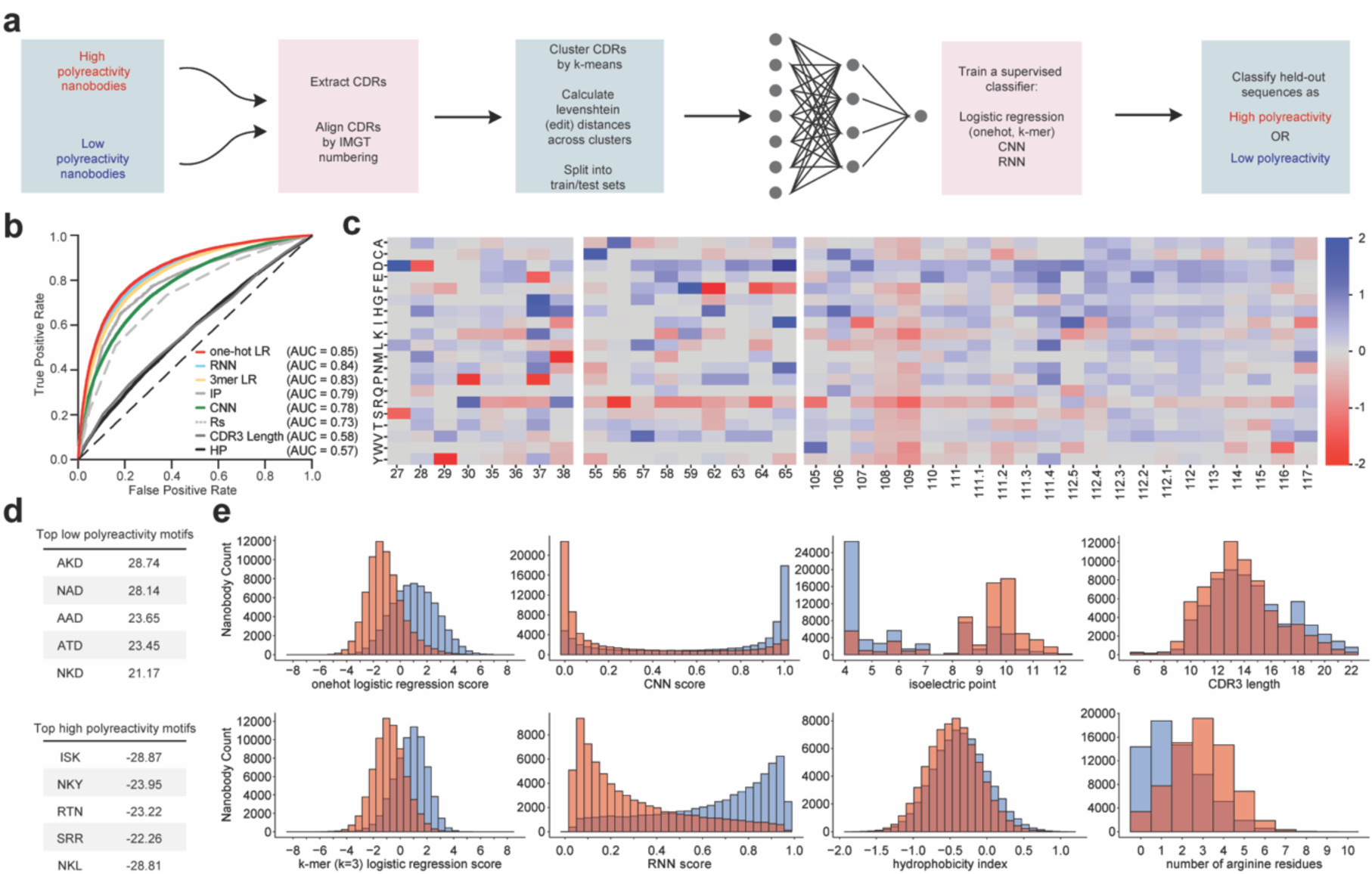
Development of computational models to predict polyreactivity. Supervised models were trained on pools of high and low polyreactivity sequences. **a**, Pipeline of computational model development, from raw NGS data to held-out predictions with sequence clustering for rigorous validation. **b**, Comparison of supervised models (one-hot and k-mer logistic regression, RNN, CNN) and biochemical properties such as hydrophobicity, isoelectric point, CDR3 lengths, and number of arginine residues. **c**, Trained parameters of a one-hot logistic regression model, showing which amino acids at specific positions are most predictive of high polyreactivity and low polyreactivity (red and blue, respectively). **d**, Polyreactivity scores of top motifs learned from a k-mer logistic regression model most predictive of low and high polyreactivity (top and bottom, respectively). **e**, Separation of high and low polyreactivity nanobodies by each of the models and biochemical properties displayed in panel b.

The one-hot logistic regression, k-mer logistic regression, and RNN models performed well at classifying distant nanobody sequences as high or low polyreacitvity, achieving 0.85, 0.83, and 0.84 Area Under Curve (AUC) respectively (Figure 3B). Whereas, the CNN (AUC=0.78, Figure 3B) achieved similar performance to metrics as described previously in literature, such as isoelectric point^16,22-24^ and the number of arginine residues^18,20,21,25^ (AUCs of 0.79 and 0.73 respectively, Figure 3B). Consistent with previous literature^15,23^, we found that hydrophobicity, as described by the hydrophobicity index, is not strongly predictive of polyreactivity (AUC of 0.57, Figure 3B). However, CDR3 length, which is a reported feature of polyreactive antibodies^16,23^ is not highly predictive of nanobody polyreactivity (AUC of 0.58, Figure 3B). Score and measurement distributions of the nanobody sequences for each of these metrics, separated by labeled class are displayed in Figure 3E.

In addition to the models’ robust performance, sequence features learned by the logistic regression methods are easily interpretable. A distinct advantage of the one-hot logistic regression model is its ability to produce a picture of amino acid contribution to polyreactivity at each position of nanobody CDR sequences (Figure 3C). In agreement with previous findings, we find that acidic residues in CDRs 2 and 3 are characteristic of low polyreactivity clones and the presence of arginine residues across all CDRs, and lysine, tryptophan, or tyrosine in CDR3 contribute to higher polyreactivity. Despite the overall enrichment of arginine and tryptophan polyreactive clones, the position specific analysis provided by the one-hot model indicates that low polyreactivity clones tolerate arginine in positions 30 and 38 of CDR1 and tryptophan in position 105 in CDR3.

Furthermore, the k-mer logistic regression model provides insight into sequence dependencies on the local level in high or low polyreactivity clones (Figure 3D). K-mer motifs containing negatively charged residues such as glutamate and aspartate are highly associated with low polyreactivity sequences, and positively charged residues such as arginine and lysine are predicted to contribute to polyreactivity, agreeing with the predictions of the one-hot logistic regression model. These motifs differ from previously reported polyreactive motifs, that were enriched in glycine and the hydrophobic amino acids valine and tryptophan^18^. However, these previously reported motifs were derived from a library where only CDR3 was diversified. We proceeded to use the one-hot and k-mer logistic regression models for further analysis based on of their accuracy and interpretability.

### Quantitative scoring of nanobody polyreactivity

In order to test if our model could go beyond predicting binary classification labels and quantitively score polyreactivity, we stained 48 nanobodies isolated from MACS and FACS pools with PSR to obtain an “index set” of sequenced clones with defined levels of polyreactivity (Figure 4A, Supplementary Table 1). Index panel nanobodies partitioned into three groups according to their level of polyreactivity: minimal polyreactivity (light gray), moderate polyreactivity (gray), and high polyreactivity (dark gray). To validate the rank order of the 48 nanobodies we measured the polyreactivity of index panel members using PSR reagent derived from solubilized HEK293 cell membranes. We found that insect cell and HEK293 derived PSR staining are highly correlated (r^2^ = 0.895, p < 0.0001), indicating that polyreactivity levels do not vary with PSR reagent type (Supplementary Figure 3C). Furthermore, to confirm that the rank order was not skewed by PSR binding to unfolded nanobodies on the surface of yeast, the index set was stained with an anti-V_HH_ antibody, which recognizes the folded nanobody framework region (Supplementary Figure 3A). Levels of anti-V_HH_ antibody staining are not correlated to insect cell PSR staining (r^2^ = 0.046, p = 0.1446, Supplementary Figure 3B), indicating that unfolded clones do not confound our dataset.

**Figure 4.**
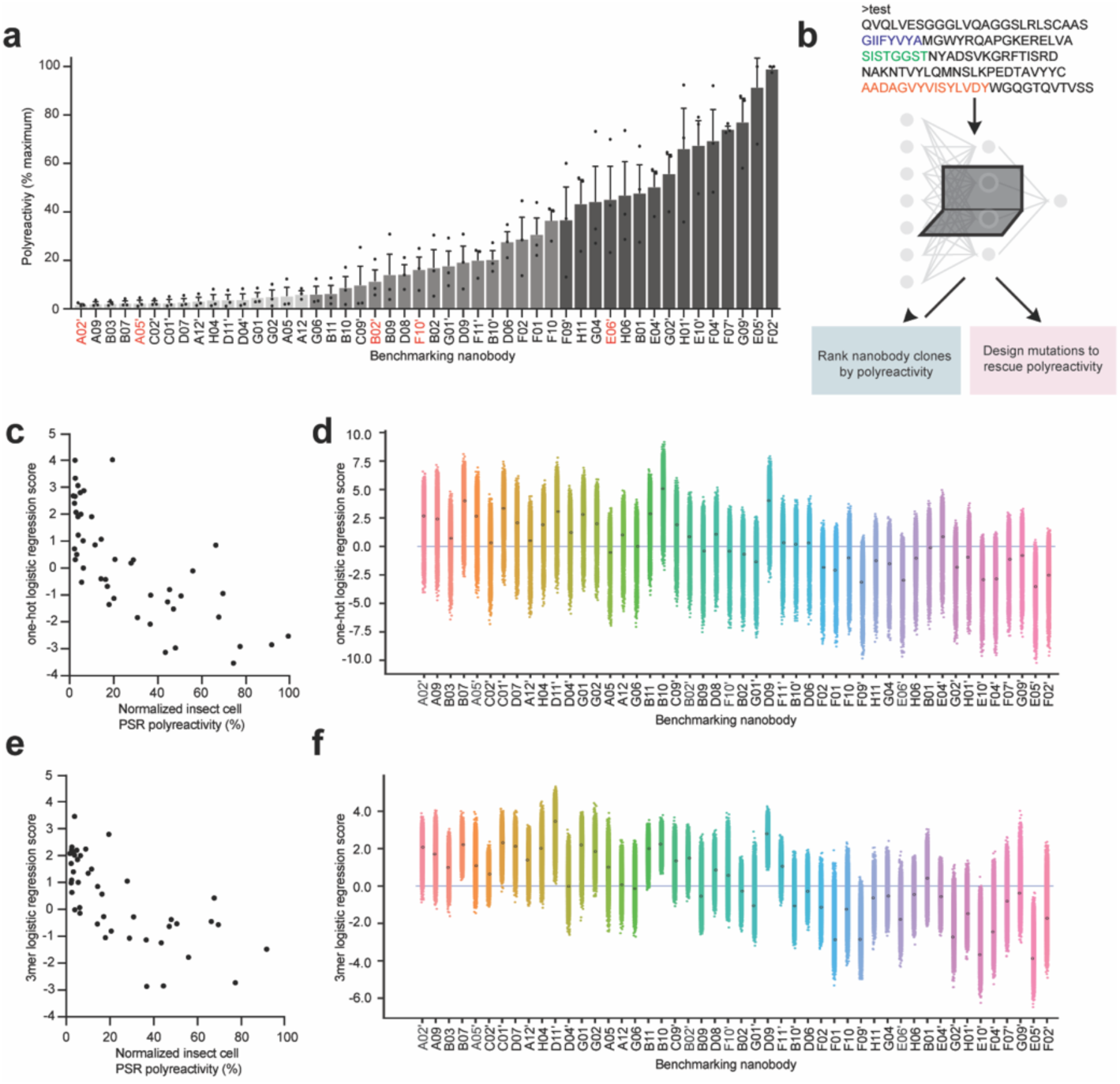
Validation of computational model for quantitative predictions of polyreactivity and design of rescue mutations. **a**, Generation of an index panel of polyreactivity mutants by *Spodoptera frugiperda* (Sf9) insect cell membranes protein polyspecificity reagent (PSR) staining of yeast displaying 48 unique nanobodies isolated from MACS enrichment as well as non-reactive and polyreactive FACS pools. Data are mean +/- SEM of three independent biological experiments performed in technical triplicate. **b**, New nanobody sequence(s) can be input into a webserver, which will output computational predictions of polyreactivity and biochemical properties of the sequence(s). It is also possible to input a nanobody sequence to retrieve top scoring rescue mutations predicted to decrease polyreactivity. **c, e**, The one-hot logistic regression model and k-mer logistic regression model trained on the full NGS dataset from FACS sorts with PSR binding were used to test quantitative predictions and rankings of the index set of clones spanning a wide range of polyreactivity levels (as measured by PSR binding) (spearman *ρ*_*s*_ of 0.77 and 0.79, respectively). **d, f**, An *in silico* double mutation scan (spanning substitutions, insertions, and deletions) was scored for predicted polyreactivity using both the one-hot logistic regression model and k-mer logistic regression model. From these *in silico* double mutation scans, a diverse set (spanning each CDR and combinations of CDRs) of high scoring mutations predicted to have low polyreactivity were selected as rescue mutations for experimental testing from two parent clones, E10’ and D06.

Biophysical characteristics of clones in our index set were reflective of the learned features in our high and low polyreactivity pools. There is a modest correlation between PSR staining of the index set and nanobody isoelectric point (r^2^ = 0.390, p < 0.0001, Supplementary Figure 3D). While nanobodies with low isoelectric points possess low polyreactivity, nanobodies with high pI values demonstrate a range of polyreactivity. Similarly, nanobody hydrophobicity index values are not correlated with polyreactivity (r^2^= 0.036, p = 0.195, Supplementary Figure 3E).

Of the 48 nanobodies, 4 were previously seen in our training set, so we did not include these in our quantitative tests. Each of the 44 remaining nanobodies had at least 6 mutations from any single nanobody sequence in the training set; the median of the minimum edit distance (a proxy for the number of mutations) of each of these index set nanobodies to the training set was 10 edit distance (the maximum similarity to the training set was 75% sequence identity). The correlation between the quantitative model predictions and the experimental binding scores to PSR, are strong - about 85% of the maximum theoretical correlation (Spearman *ρ*_*s*_ of 0.77 and 0.79, for the one-hot and k-mer logistic regression models, respectively) (Figure 3B). For comparison, the Spearman correlations between the three independent biological replicate experiments were 0.87, 0.87, and 0.95. Thus, our models trained on sequence pools of high and low polyreactivity nanobody CDR sequences are highly accurate for both classification and regression tasks for clones with distinct sequences.

### Model performance at predicting polyreactivity of closely related sequences

To determine if our computational model could accurately assess the influence of point mutations in single nanobody clones, we utilized the autonomous hypermutation yeast surface display (AHEAD) error-prone DNA replication system^32^ to rapidly evolve the four most polyreactive clones from our index set (Nb E05’, F02’, G09’, and F07’) to have reduced binding to the PSR reagent. Over the course of four AHEAD cycles involving nanobody hypermutation and FACS sorting, global PSR staining of the evolved nanobody population decreased (Supplementary Figure 4). Deep sequencing analysis following the fourth FACS round revealed variation in the CDR regions of each of the four nanobodies.

A large proportion of the clones enriched by AHEAD are predicted to have reduced polyreactivity by both the one-hot and 3-mer logistic regression models. For the four clones, 97%, 67%, 69%, and 93% of the observed mutations are predicted to decrease polyreactivity by the one-hot logistic regression model, with similar decreases predicted by the k-mer logistic regression model (Supplementary Table 2). Furthermore, K31E^36^, A50T^55^, and R57P^64^ substitutions that arose in nanobody E05’ reflect the position specific analysis provided by the one-hot logistic regression model, where K, R, and A are characteristic of polyreactive nanobodies at positions 36, 55, and 64 and all three substitutions are characteristic of clones with reduced polyreactivity (Figure 3C). In a computational ranking of the polyreactivity of all 494 single amino acid substitutions using the one-hot logistic regression model in the CDR regions of E05’ found in our AHEAD experiment, from lowest to highest, R57P^64^ ranked 28^th^, K31E^36^ ranked 37^th^, and A50T^55^ is 101^st^. Overall, the AHEAD-based directed evolution experiment produces clones that our computational models predict to have reduced polyreactivity suggesting that our models can accurately score the polyreactivity of closely related sequences.

With confidence in our models’ performance on related clones, we employed our computational model to independently predict sequence substitutions to reduce polyreactivity of the highly polyreactive clone E10’ and moderately polyreactive clone D06 from our index set. We performed a comprehensive *in silico* single and double mutant scan, scored each sequence with both the one-hot logistic regression model and the k-mer logistic regression model (Figure 4B-D), and ranked all the possible single and double mutants, including insertions and deletions, surrounding the seed sequence. We sampled the substitutions most likely to reduce polyreactivity (with the exception of a substitution that would have introduced a cysteine that could disrupt disulfide bond formation) by selecting diverse mutations across residue types and positions that are contained within a single CDR and span each of the possible combinations of different CDR regions. Furthermore, if there was a mutation indicated to decrease polyreactivity by the k-mer logistic regression that scored similarly according to the one-hot logistic regression model, we selected the sequence with a higher k-mer logistic regression score to take into account local sequence dependencies. We selected the three top scoring single mutations for each of the CDR regions, the top scoring double mutants within a single CDR region, and the top scoring double mutants spanning two CDR regions where at least one of the individual single mutations had not already been tested in a different combination.

For the moderately polyreactive D06 nanobody, 18 out of 21 variants that were computationally designed to decrease polyreactivity reduced levels of binding to insect cell PSR staining (Figure 5A). More stringently, 11 out of 21 mutations exhibited at least two-fold reductions in polyreactivity. Although substitutions in each of the CDR regions were able to lower polyreactivity, CDR3 appeared to drive polyreactivity as the most significant reductions in polyreactivity occurred from variations in the CDR3 region including A97H^106^ and R98D^107^ R99H^108^.

**Figure 5.**
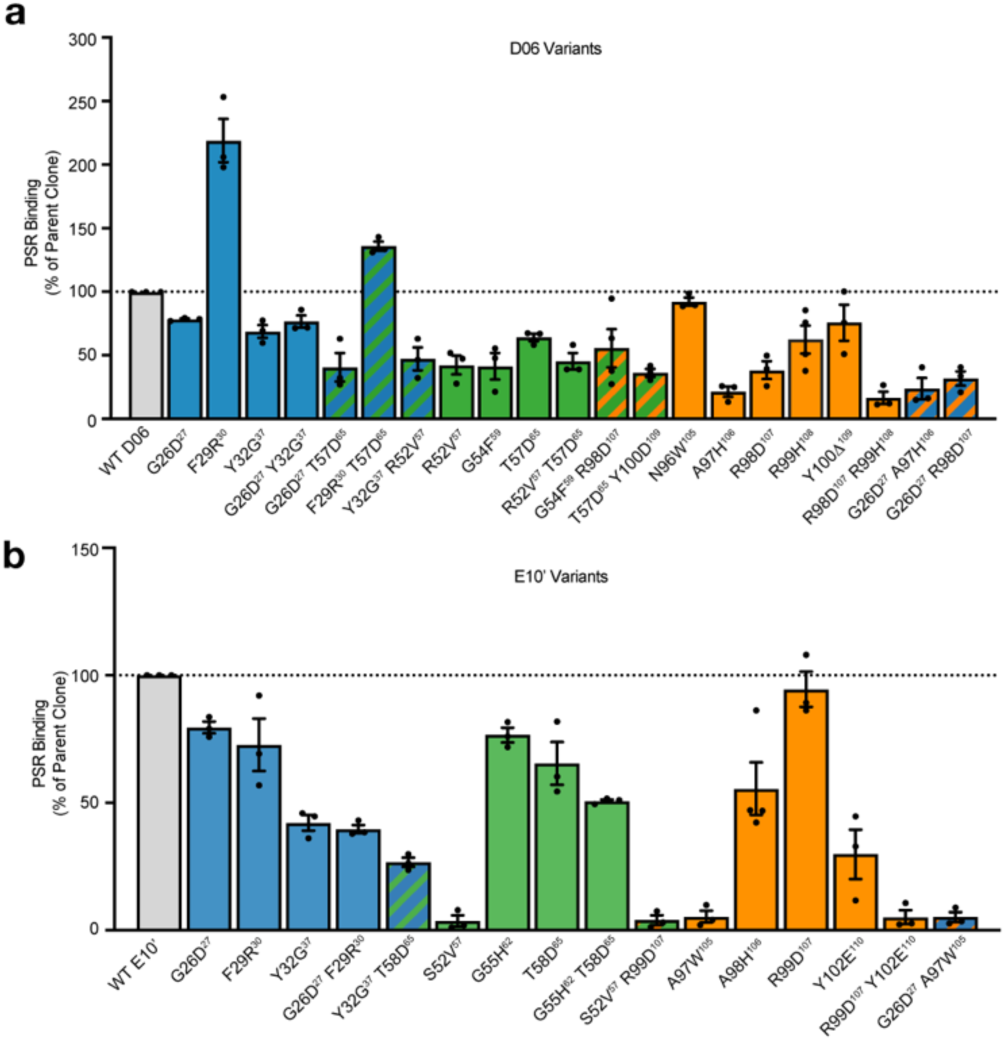
*In silico* designed substutions reduce nanobody polyreactivity. **a**, Polyspecificity reagent (PSR) staining of yeast displaying D06 variants. For the moderately polyreactive D06 nanobody, 18 out of 21 variants that were computationally designed to decrease polyreactivity reduced levels of binding to insect cell PSR staining Data in **a** comprise the mean +/- SEM of at least three independent experiments, each performed in technical triplicate. **b**, PSR staining of yeast displaying E10’ variants. For the highly polyreactive E10’ nanobody, 15 out of 16 computationally predicted single and double substitutions reduced binding to PSR reagent. Data in **b** comprise the mean +/- SEM of at least three independent experiments, each performed in technical triplicate.

For the highly polyreactive E10’ nanobody, 15 out of 16 computationally predicted single and double substitutions reduced binding to PSR reagent (Figure 5B). 9 out of the 16 substitutions reduced polyreactivity by at least 50%, including mutations in each of the three CDR regions. Strikingly, the R99D^107^ Y102E^110^ clone, which was predicted to have the lowest polyreactivity value using the k-mer logistic regression model has very low polyreactivity by experimental PSR staining.

### Reducing polyreactivity of a functional clone

We next tested if our model could be employed to decrease the polyreactivity of nanobody clone that was independently selected for antigen specificity. AT118i4h32 is a nanobody antagonist for the angiotensin II type 1 receptor (AT1R), a G protein-coupled receptor (GPCR) that is a central regulator of blood pressure and renal function. AT118i4h32 directly competes with the binding of small molecule and peptide ligands to the AT1R and is active *in vivo*, reducing mouse blood pressure in a comparable degree to the clinically used angiotensin receptor blocker losartan^28^. Additionally, AT118i4h32 has been humanized with 11 amino acid substitutions to resemble a human V_H_3-23. Although pharmacologically intriguing, AT118i4h32 is highly polyreactive in the PSR assay and has a high pI value (9.6), which is characteristic of polyreactive antibodies. Furthermore, a crystal structure of AT118i4h32 displays large patches of positive charge on the protein surface (Figure 6a, Supplementary Table 3) and enrichment of both solvent exposed arginine and hydrophobic residues in the CDR regions (Figure

**Figure 6.**
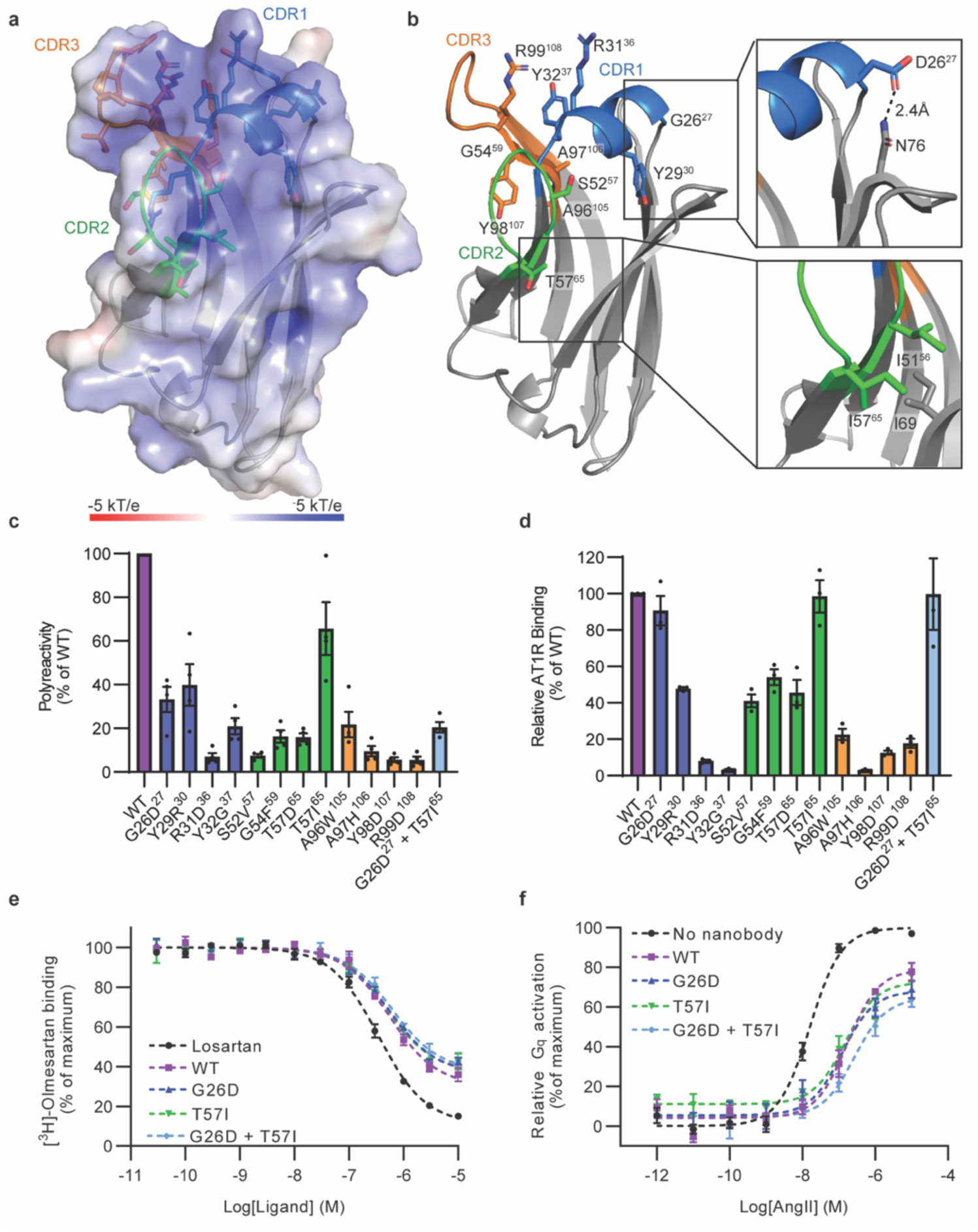
Development of AT118i4h32 variants with reduced polyspecificity. **a**, electrostatic surface of AT118i4h32. CDR1, CDR2, and CDR3 are colored blue, green, and orange. All positions substituted to produce variants of AT118i4h32 with reduced polyreactivity are shown in sticks with atomic coloring **b**, AT118i4h32 structure as colored in a. G26D^27^ and T57I^65^ substitutions are boxed. **c**, PSR staining of yeast displaying AT118i4h32 variants. All amino acid substitutions decrease polyreactivity. Data in c comprise the mean +/- SEM of four independent experiments, each performed in technical triplicate. **d**, binding of AT118i4h32 variants to HEK293 suspension cells expressing FLAG-AT1R. Cells were stained with AT118i4h32-V5-His variants, AlexaFlour-488 conjugated anti-FLAG, and AlexaFlour-647 conjugated anti-V5 antibodies, then analyzed by flow cytometry. Data in d is the average of three independent experiments performed in technical triplicate, error bars are shown as SEM. **e**, radioligand competition binding of AT118i4h32 variants or the small molecule antagonist losartan and [^3^H]-olmesartan to AT1R in cell membranes. Like WT AT118i4h32, the G26D, T57I, and G26D+T57 variants compete with olmesartan for binding to the AT1R. Data in e is the average of three independent experiments performed in technical triplicate, error bars are shown as SEM. **f**, suppression of Gq-mediated inositol monophosphate production by AT118i4 in response to AngII stimulation. HEK293 suspension cells expressing FLAG-AT1R were treated with 5 μM AT118i4h32 or no nanobody prior to AngII stimulation. Data in d is the average of three independent experiments performed in technical triplicate, error bars are shown as SEM. K_>i_ values are reported in Supplementary Table 3.

We analyzed the sequence of AT118i4h32 and selected twelve single amino acid substitutions scattered throughout each CDR predicted to reduce polyreactivity based on the one-hot logistic regression model. AT118i4h32 variants were displayed on the surface of yeast and all showed reduced levels of PSR binding (Figure 6C). Neutralizing the highly basic patch composed of R30^35^, R31^36^, and R99^108^ on the surface of AT118i4h32 (Figure 6A) with R31D^36^ and R99D^108^ substitutions substantially reduces AT118i4h32 polyreactivity. Notably, introduction of an additional arginine residue with the Y29R^30^ substitution, which introduces a RRR sequence motif into CDR1, reduces polyreactivity, further demonstrating that arginine’s contribution to polyreactivity is highly position dependent.

To assess the effects of these substitutions on antigen binding, AT118i4h32 variants were recombinantly expressed in *E. coli* and purified to evaluate AT1R binding by flow cytometry (Figure 6D). Two AT118i4h32 variants, G26D^27^ and T57I^65^, retained at least 80% of wild-type binding levels to the AT1R. Combination of the G26D^27^ and T57I^65^ substitutions retained high levels of binding to the AT1R and yielded a clone with a modest decrease in PSR binding compared to the G26D^27^ variant (Figure 6C), bringing the overall level of polyreactivity close to that of the clinically approved nanobody drug Cablivi/caplacizumab^33^ (Supplementary Figure 5A). Additionally, the G26D^27^, T57I^65^ variant has reduced polyreactivity compared to the wild-type nanobody as measured by ELISA assay (Supplementary Figure 5B-G). AT118i4h32 variants containing G26D^27^ and T57I^65^ maintain the ability to act as receptor antagonists, displacing small molecule orthosteric antagonists (Figure 6E) and suppressing receptor signaling upon angiotensin II (AngII) stimulation (Figure 6F).

To investigate how the G26D^27^ T57I^65^ substitutions alter AT118i4h32’s structure and contribute to reduce polyreactivity, we crystallized AT118i4h32 G26D^27^ T57I^65^ and solved the structure at 1.6 Å resolution (Figure 6B, Supplementary Table 3). The T57I^65^ substitution is located at the end of CDR2. I57^65^ forms more favorable hydrophobic interactions with neighboring I51^56^ and I65 side chains than T57^65^. In the case of AT118i4h32, maintaining this hydrophobic interaction is essential for antigen recognition, as the T57D^65^ substitution diminished AT1R binding two-fold (Figure 6D). While the T57I^65^ mildly decreases polyreactivity, AT118i4h32 variants containing the T57I^65^ substitutions had slightly decreased thermal stability (Supplementary Table 4), indicating that changes in reduced polyreactivity are not necessarily correlated with thermal stability.

Residue D26^27^, found at the N-terminus of helical CDR1, forms a hydrogen bond with the side chain of framework residue N76 in all eight copies of the nanobody in the crystal structure’s asymmetric unit (Figure 6B). This hydrogen bond rigidifies the CDR1 position and may reduce the flexibility of the nanobody’s CDR regions. Additionally, the G26D substitution improves AT118i4h32’s stability; we observed a five-fold increase in AT118i4h32 G26D^27^ yield from *E. coli* and a two degree increase in melting temperature of the G26D^27^ variant (Supplementary Table 4) over wild-type levels. Corresponding G26D^27^ substitutions reduced the polyreactivity of nanobodies D06 and E10’. Despite occurring in just 0.05% of sequences from the naïve repertoire of seven llamas^34^ (1.12 million unique nanobody sequences), the D27 substitution may be both beneficial and tolerated in many sequence contexts and may broadly reduce polyreactivity by reducing the conformational flexibility of the CDR regions^35^.

### Expansion of computational method

Upon examination of corresponding substituted positions in D06, E10’, and AT118i4h32 we observe some substitutions reduce polyreactivity in all clones, such as G26D^27^, whereas other mutations dramatically reduced polyreactivity of some nanobodies (i.e., E10’ A97W^105^ and AT118i4h32 A96W^105^) while having little to no effect in another clone (i.e., D06 N96W^105^). This suggests that *position dependency is critical for polyreactivity*, which may be more accurately captured with a larger data set. Therefore, we sought to improve our *in silico* method with expanded sequencing data. Through additional rounds of FACS selection, we collected 1,221,800 unique low polyreactivity clones and 1,058,842 unique high polyreactivity clones. We trained our suite of supervised classification models on this extended dataset and included analysis of an extra position at the end of CDR2, which has some variability in the synthetic nanobody library, but was not included in the initial analysis.

To test classification accuracy, we clustered the sequences into 10 clusters using a k-means algorithm for train/test splits, and again limited our training dataset to sequences with at least 10 mutations as compared to any sequence in the test sets. We achieved comparable classification AUCs to the logistic regression and RNN models trained on the original FACS sorts (one-hot logistic regression: 0.83, 3-mer logistic regression: 0.83, RNN: 0.84) (Supplementary Figure 6A). The convolutional neural network model received a significant performance boost (CNN: 0.83 compared to previously 0.78 AUC) (Supplementary Figure 6A). For the higher throughput dataset, we see that the models that capture more complexities in sequences, such as the CNN and RNN, have higher accuracies, suggesting that there are meaningful dependencies in nanobody sequences that contribute to polyreactivity beyond site-specific amino acid contributions and/or 3-mer motifs and would allow us to make more accurate predictions to reduce polyreactivity for individual sequences. Furthermore, for each of these models we see an improved correlation (Spearman R) of polyreactivity scores with the index set measurements (one-hot logistic regression: 0.87, 3-mer logistic regression: 0.86, CNN: 0.88, RNN: 0.88) (Supplementary Figure 6B-E). The majority of substitutions applied to clones D06, E10’, and AT118i4h32 are still predicted to decrease polyreactivity across the four models trained on the deeper FACS sequencing experiments (37, 37, 41, and 23 out of 45 mutations for one-hot logistic regression, k-mer logistic regression, CNN, and RNN respectively; for the RNN in particular, most mutations that were not predicted to decrease polyreactivity had very small changes in predicted signal, Supplementary Table 6).

As a resource to the field, we provide open-access use of our polyreactivity prediction software on our webpage (http://18.224.60.30:3000/). The webserver allows users to input a nanobody sequence(s) in FASTA format and outputs the aligned nanobody sequence with IMGT numbering using ANARCI^30^, along with biochemical properties of the sequence, including isoelectric point, hydrophobicity, CDR definitions (IMGT), CDR lengths, and computational predictions of polyreactivity scores using the one-hot logistic regression models that were trained for the design of rescue mutations.

## DISCUSSION

Previous work has identified some biophysical characteristics of polyreactivity, but these studies have generally been performed on relatively small sets of antibody sequences without an explicit attempt to improve polyreactivity properties. Here, we designed and conducted high-throughput experiments to capture diverse clones that were not influenced by other selection pressures, facilitating an unbiased analysis of nanobody polyreactivity. Starting with a large naïve synthetic library mimicking the llama immunological repertoire, we isolated large pools of high and low polyreactivity nanobody clones based upon binding to the mixed-protein PSR reagent. Our models are over 80% accurate in discriminating between clones with high and low polyreactivity (Figure 3B), rank levels of polyreactivity with high fidelity (Figure 4), and reliably identify amino acid substitutions that reduce polyreactivity (Figures 5 and 6C).

Since our models were built upon experiments that were intentionally designed to interrogate sequence contributions to polyreactivity, they are highly accurate at measuring polyreactivity. In accordance with previous studies, our deep dive results suggest that arginine generally promotes nanobody polyreactivity while glutamate and asparate usually decrease polyreactivity. However, we find amino acid contributions to polyreactivity are highly position dependent and more nuanced than broad generalizations suggest. This finding is in agreement with a recent independent study that analyzed polyreactivity of a subset of antibodies^17^. Furthermore, our computational models’ ability to accurately quantify polyreactivity from sequence identity constitutes a large step forward as we can diagnose and engineer away polyreactivity of existing clones. More complex models including the CNN and RNN models also allowed us to evaluate dependencies of amino acids in different locations in nanobodies to polyreactivity. We find such dependencies contribute to polyreactivity indicating that both local and global characteristics of nanobodies influence their degree of polyreactivity.

We provide to the community an easy-to-use webserver that encapsulates our computational methods. These methods can guide antibody discovery campaigns at many points in the discovery pipeline. For instance, our software can be used to prospectively predict amino acid substitutions that will reduce polyreactivity of a single clone such as AT118i4h32. Moreover, the polyreactivity of a list of antigen binders can be ranked for clone prioritization during selection campaigns. We found that substitutions in each of the CDR regions of D06, E10’, and AT118 reduce polyreactivity, suggesting that each CDR region contributes to polyreactivity. Therefore, if a certain CDR region is critical for antigen recognition, substitutions in alternative CDR regions can potentially compensate in reducing polyreactivity. In addition, our success in reducing polyreactivity of AT118i4h32, where the humanized framework region differs from clones in the training set, indicates that our methods are applicable to nanobodies from a range of sources. Although outside the scope of this manuscript, similar approaches can be applied to conventional antibodies, adding in the three light-chain CDRs and germline gene choice as additional factors for polyreactivity prediction and optimization.

### Statistical Methods

Prism software (Graphpad) was used to analyze data and perform error calculations. Data are expressed as arithmetic / geometric mean ± SEM or arithmetic / geometric mean ± SD.

### Data Code Availability Statement

The code for scoring new sequences for polyreactivity, designing rescue mutations, training polyreactivity models, and calculating biochemical properties of a sequence can be found on github: https://github.com/debbiemarkslab/nanobody-polyreactivity, and the webserver is available here: (http://18.224.60.30:3000/). Coordinates and structure factors for the AT118i4h32 structures are deposited in the Protein Data Bank under accession codes 7T83 and 7T84.

## ACKNOWLEDGMENTS

This work was funded by a Merck Postdoctoral Fellowship from the Helen Hay Whitney Foundation to M.A.S.; NIH training grant 5T32GM132089-03 to V.G.M; NIH TR01 grant 1R01CA260415 to C.C.L, D.S.M., and A.C.K; 5R21HD101596 to A.C.K.; the Moore Inventor Fellowship to C.C.L. We thank Dr. Laura Wingler and Dr. Dean Staus for providing AT118i4h32 for crystallization experiments and Dr. Marie Bao for critical reading of the manuscript. We thank the staff at Advanced Photon Source GM/CA beamlines for support of X-ray data collection. GM/CA@APS is funded by the National Cancer Institute (ACB-12002) and the National Institute of General Medical Sciences (AGM-12006, P30GM138396). The Eiger 16M detector at GM/CA-XSD was funded by NIH grant S10 OD012289. Portions of this research was conducted at the Advanced Photon Source, a U.S. Department of Energy (DOE) Office of Science User Facility operated for the DOE Office of Science by Argonne National Laboratory under Contract No. DE-AC02-06CH11357. We thank SBGrid Consortium for structural biology software support. D.S.F. experiments were carried out at the Center for Macromolecular Interactions in the Department of Biological Chemistry and Molecular Pharmacology at Harvard Medical School with support from Dr. Kelly Arnett.

## AUTHOR CONTRIBUTIONS

M.A.S., E.P.H., J.S., D.S.M., A.C.K designed research. M.A.S. and E.P.H. performed MACS and FACS selections. E.P.H., M.A.S., and G.R.N. analyzed nanobody polyreactivity. J.S. and A.Y.S. designed computational algorithm. A.W. performed AHEAD experiments under the supervision of C.C.L. J.S. and E.P.H. analyzed AHEAD evolution experiments. G.R.N., J.H., E.P.H. and M.A.S purified nanobody variants. E.P.H. and J.H. performed nanobody size exclusion chromatography, E.P.H., J.H., and V.M. developed and ran ELISA assays. E.P.H. and J.H. performed and analyzed anti-nanobody antibody staining experiments. J.K.M. and A.Y.S. designed webserver. M.A.S. generated PSR reagent, performed mammalian cell binding, thermal stability, radioligand binding, and AT1R signaling assays. M.A.S. and G.R.N. determined the crystal structures of AT118i4h32. M.A.S., E.P.H., and J.S. wrote the manuscript with input from all authors.

## COMPETING INTERESTS STATEMENT

C.C.L is a co-founder of K2 Biotechnologies Inc., which applies continuous evolution technologies to antibody engineering. D.S.M. is an advisor for Dyno Therapeutics, Octant, Jura Bio, Tectonic Therapeutic and Genetech, and is a co-founder of Seismic Therapeutic. A.C.K. is a co-founder and consultant for Tectonic Therapeutic and Seismic Therapeutic and for the Institute for Protein Innovation, a non-profit research institute.

