## Supplementary_information for "An *in silico* method to assess antibody fragment polyreactivity"

Supplementary Figures

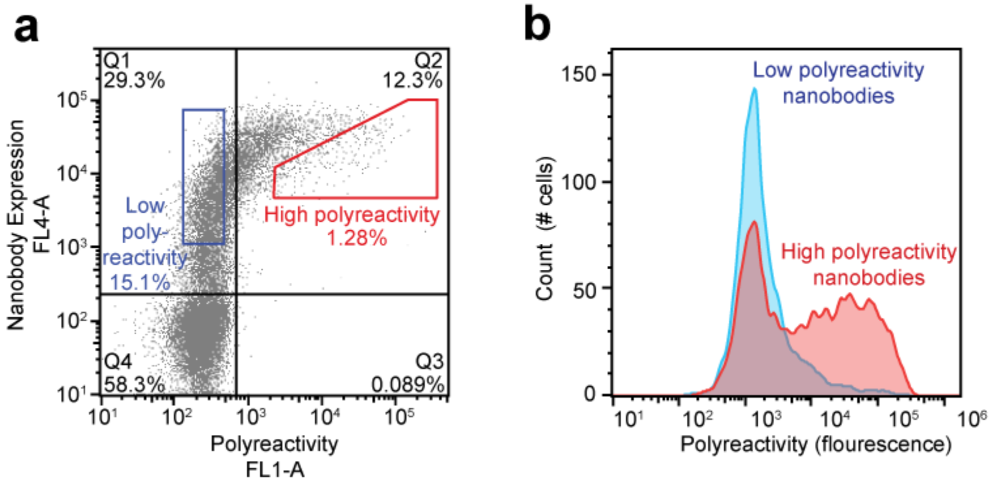


**Supplementary Figure 1. FACS Gating strategies. a**, the nanobody library enriched for PSR binders via MACS was stained with biotinylated insect cell PSR reagent, followed by AlexaFlour-647 conjugated αHA-antibody to assess nanobody expression on the yeast cell surface and AlexaFlour-488 conjugated streptavidin. Highly expressing yeast representing the low polyreactivity (PSR negative) and high polyreactivity (PSR positive) populations were collected. **b**, Analytical flow cytometry staining illustrates the difference in polyreactivity between the low polyrectivity and high polyreactivity nanobody pools.

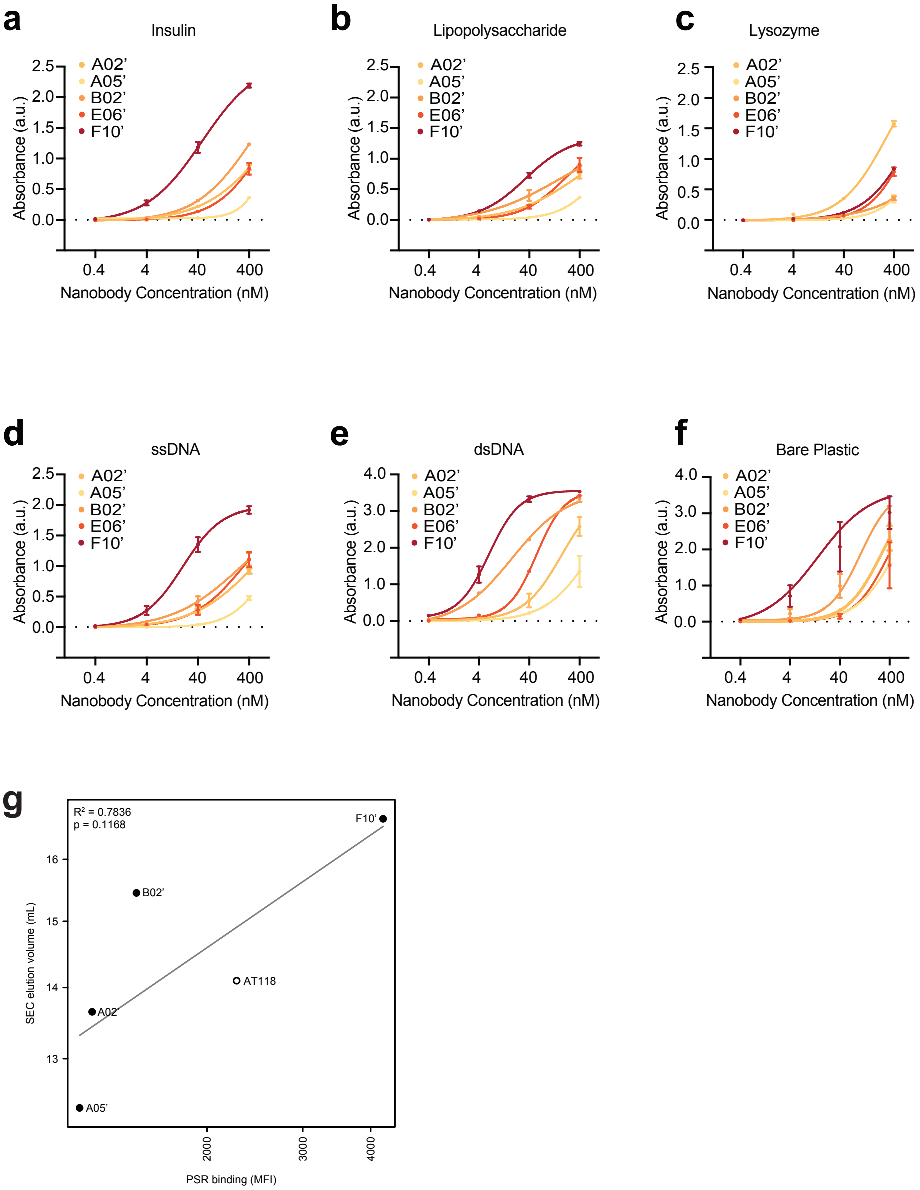


**Supplementary Figure 2. a-f,** Polyreactivity of index panel members as measured by nanobody binding to the indicated reagents in direct ELISAs. Data are mean +/- SEM of two independent experiments, each performed with technical triplicates. **g,** Correlation between nanobody size exclusion chromatography (SEC) elution volume and PSR staining, indicating that nanobodies with higher polyreactivity tend to elute later than less polyreactive nanobodies.

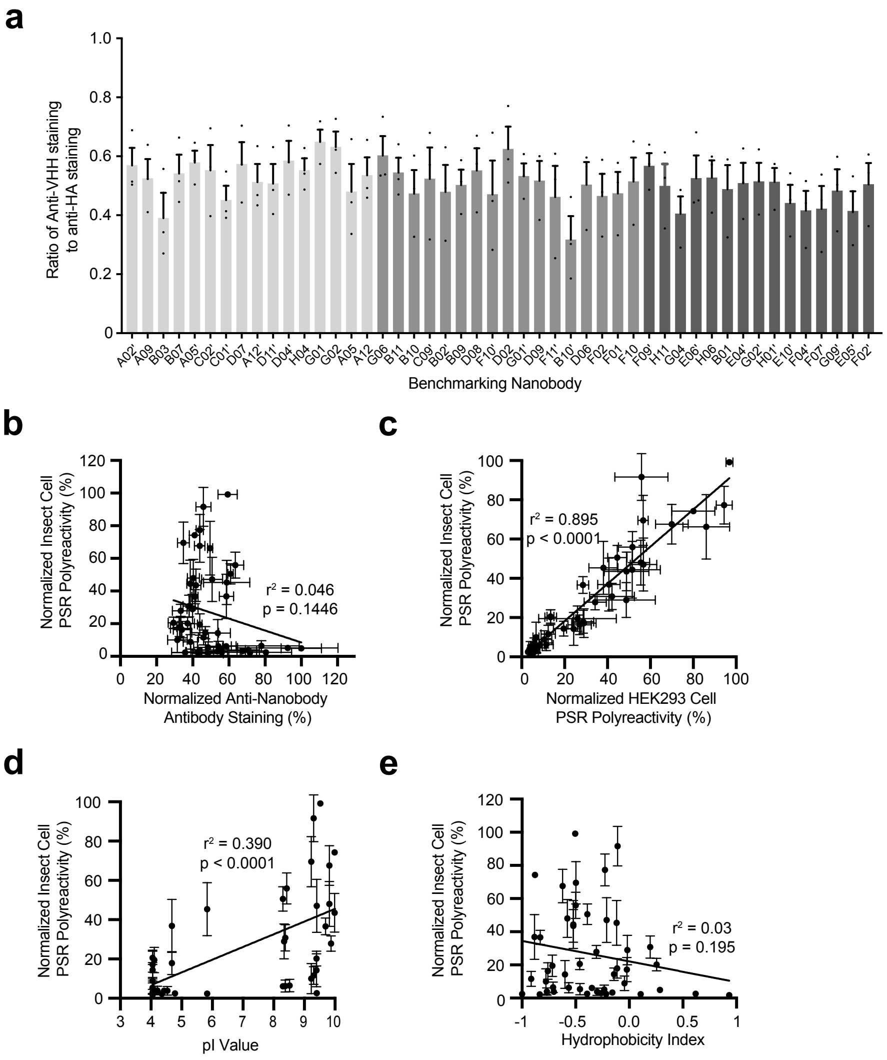


**Supplementary Figure 3.** **a,** Anti-nanobody antibody staining of yeast displaying polyreactivity index panel members, indicating that all panel members are folded. Data are mean +/- SEM of three independent biological experiments performed in technical triplicate. **b,** anti-nanobody antibody staining is not correlated with insect cell PSR staining ($r^2 = 0.046$). **c,** Insect cell PSR staining of index panel members is well correlated with HEK293 cell PSR staining, suggesting that polyreactive panel members are not binding to specific proteins in insect cell membranes. HEK293 cell PSR staining data are mean +/- SEM of three independent biological experiments. **d,** Correlation between index panel nanobody pI values and insect cell PSR reagents. Nanobodies with low pI values tend to possess low polyreactive, while nanobodies with high pI values exhibit a range of polyreactivity. **e,** Index panel nanobody hydrophobicity index values and insect cell PSR reagent staining are not correlated.

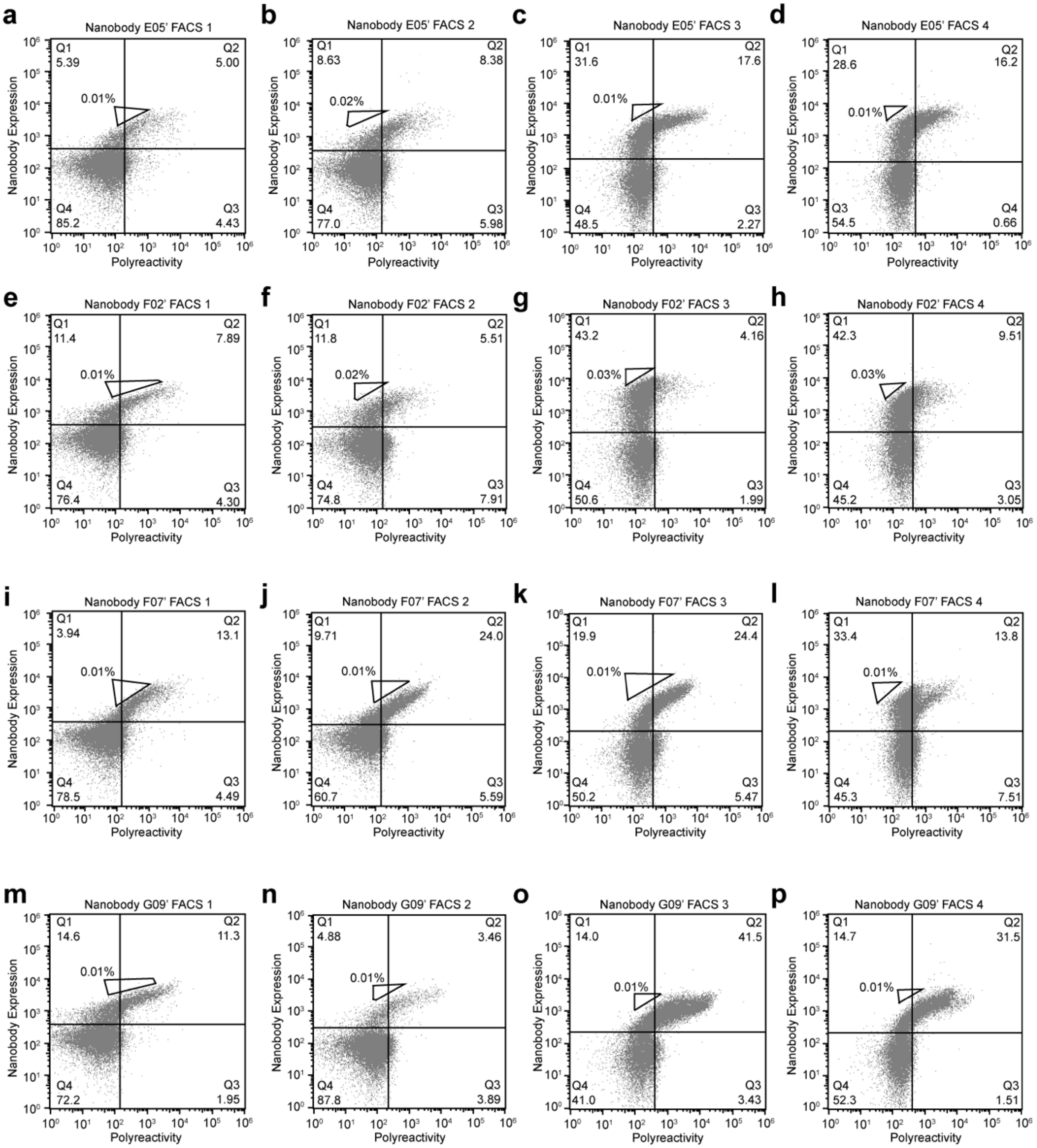


**Supplementary Figure 4.** Evolution of index panel nanobodies using the AHEAD system to reduce polyreactivity. **a-d,** Selected FACS plots showing polyreactivity reduction of nanobody E05'. **e-h,** Selected FACS plots showing polyreactivity reduction of nanobody F02'. **i-l,** Selected FACS plots showing polyreactivity reduction of nanobody F07'. **m-p,** Selected FACS plots showing polyreactivity reduction of nanobody G09'.

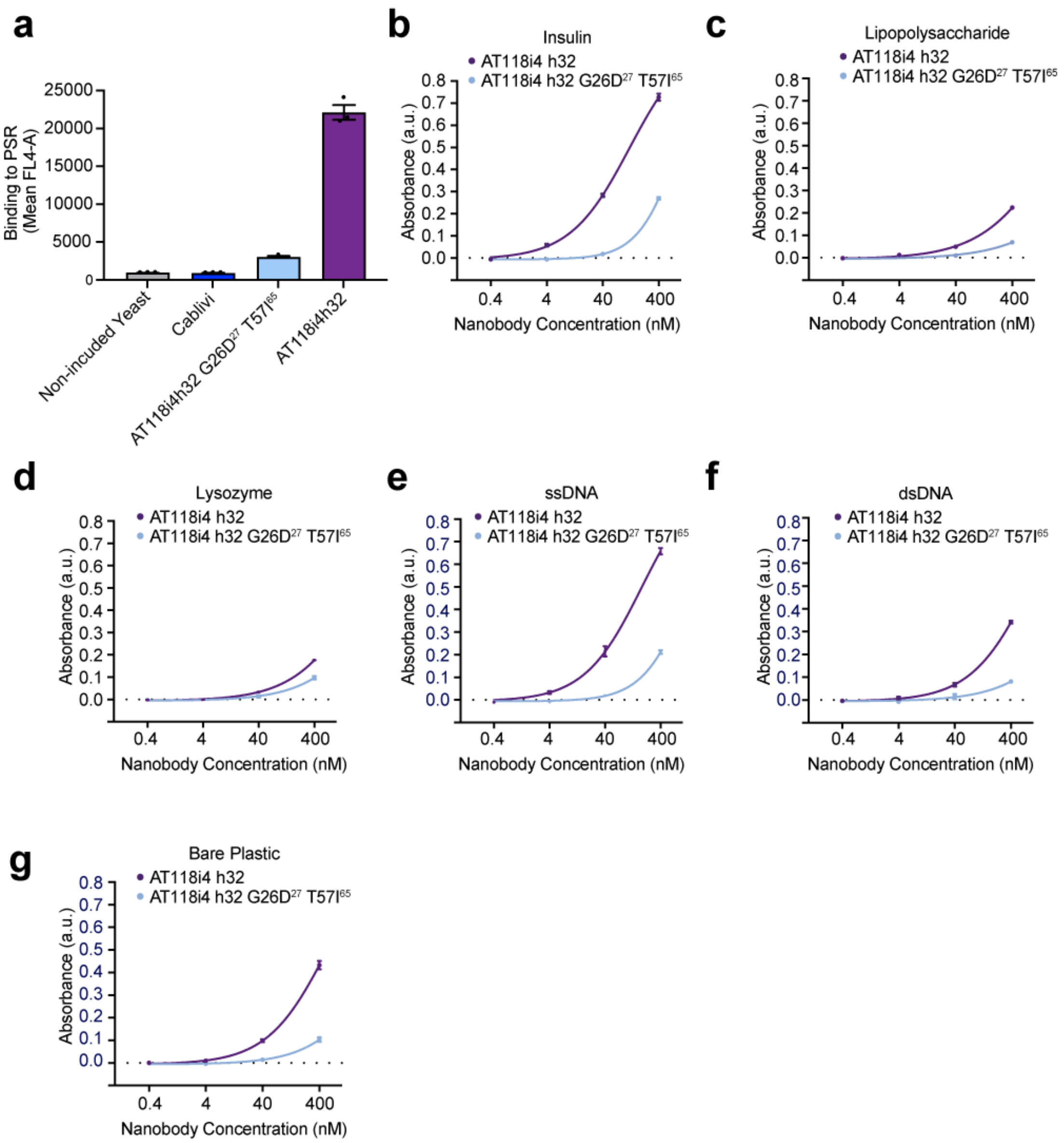


**Supplementary Figure 5 a,** Polyreactivity of non-induced yeast not expressing nanobody, AT118i4h32, AT118i4 h32 G26D$^{27}$ T57I$^{65}$, and the clinically approved nanobody drug Cablivi, as measured by insect cell PSR staining. Data is representative of three independent experiments. Error bars represent SEM of three technical replicates. **b-g,** Polyreactivity of AT118i4 h32 and AT118i4 h32 G26D$^{27}$ T57I$^{65}$, as measured by binding of AT118 variants to the specified reagents by direct ELISA assays. Data are mean +/- SEM and are representative of two independent experiments, each performed with technical triplicates.

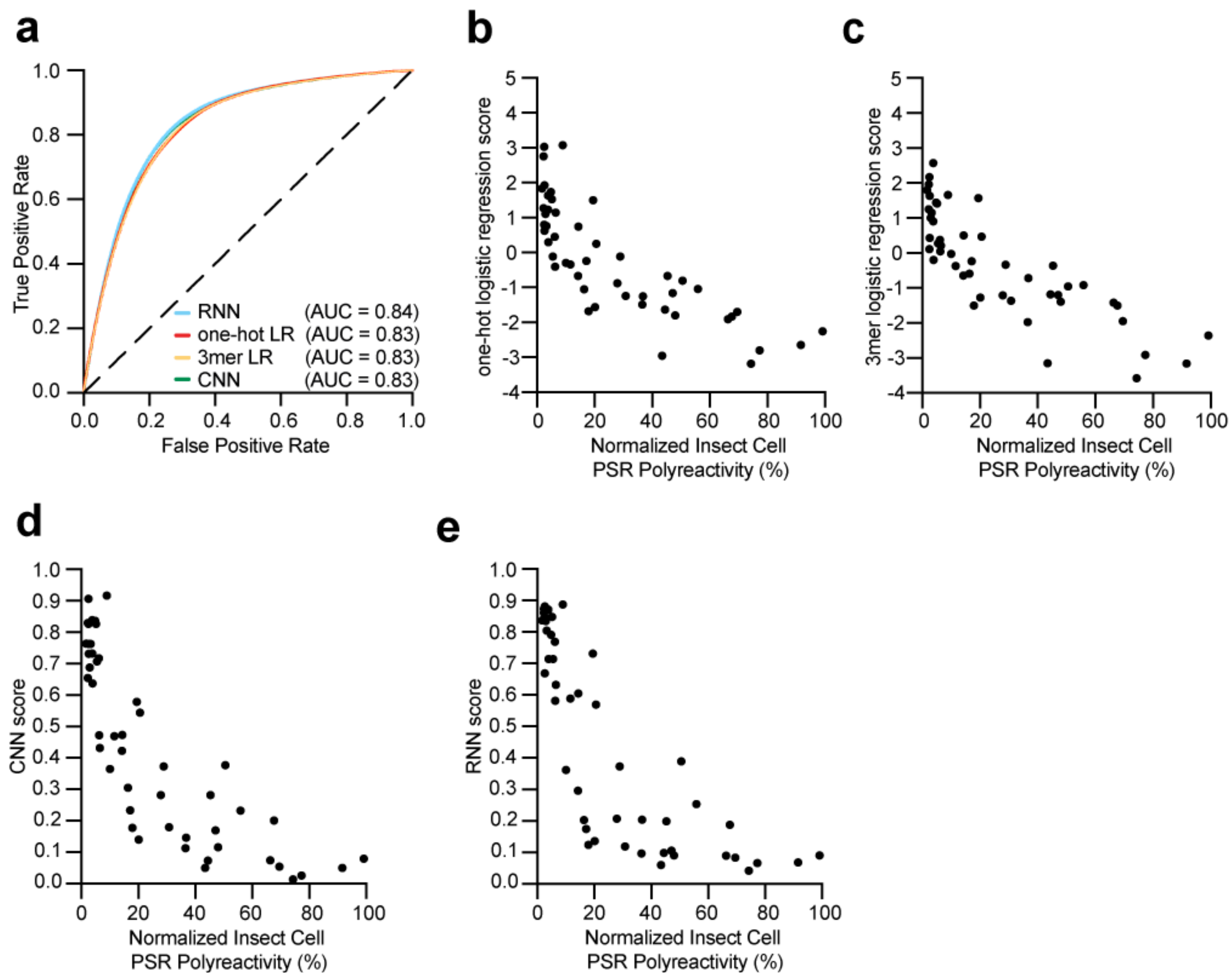


**Supplementary Figure 6. a,** Comparison of supervised models (one-hot and k-mer logistic regression, RNN, CNN) trained on a deeper FACS sort of millions of high and low polyreactive sequences. Sequences were clustered into ten validation clusters and models were trained in cross-fold validation using only sequences greater than 10 mutations away from any test sequence in the training set. **b-e,** Quantitative predictions of polyreactive compared to the experimentally measured PSR levels of the index set of clones. The one-hot logistic regression, k-mer logistic regression, CNN, and RNN models were trained on the full NGS dataset (except for sequences that had the exact same CDR sequences as sequences in the index set) and were used to predict relative polyreactivity (spearman $\rho_s$ of 0.87, 0.86, 0.88, 0.88) respectively.

### Supplementary Tables

### Table S1. Sequences of nanobodies

| **Name** | *Italics CDR Region*<br>**Sequence (Bold = mutation from corresponding parent; only non-synonymous mutations are indicated)** |
|---|---|
| A02’ | QVQLVESGGGLVQAGGSLRLSCAAS*GIIFYVYA*MGWYRQAPGKE<br>RELVA*SISTGGST*NYADSVKGRFTISRDNAKNTVYLQMNSLKPE<br>DTAVYYC*AADAGVYVISYLVDY*WGQGTQVTVSS |
| C01’ | QVQLVESGGGLVQAGGSLRLSCAAS*GSIFLLYA*MGWYRQAPGKE<br>RELVA*AITIGGST*NYADSVKGRFTISRDNAKNTVYLQMNSLKPE<br>DTAVYYC*NAPEVIRPYFVAYDY*WGQGTQVTVSS |
| A05’ | QVQLVESGGGLVQAGGSLRLSCAAS*GSIFRRNA*MGWYRQAPGKE<br>RELVA*RISWSGGST*YYADSVKGRFTISRDNAKNTVYLQMNSLKP<br>EDTAVYYC*NADNEDYLKVQSDMDY*WGQGTQVTVSS |
| A09 | QVQLVESGGGLVQAGGSLRLSCAAS*GKYNA*MGWYRQAPGKEREF<br>VA*AISLGGSTT*YYADSVKGRFTISRDNAKNTVYLQMNSLKPEDT<br>AVYYC*ARDGPYDAHSLEDDY*WGQGTQVTVSS |
| B07 | QVQLVESGGGLVQAGGSLRLSCAAS*GFIFHYYV*MGWYRQAPGKE<br>RELVA*AITSDDS*TYADSVKGRFTISRDNAKNTVYLQMNSLKPED<br>TAVYYC*AVDVWYSTDHFSDYDY*WGQGTQVTVSS |
| H04 | QVQLVESGGGLVQAGGSLRLSCAAS*GFIFRYNA*MGWYRQAPGKE<br>RELVA*TITSSDGST*YYADSVKGRFTISRDNAKNTVYLQMNSLKP<br>EDTAVYYC*AEFWVGVYNHPGYDY*WGQGTQVTVSS |
| D07 | QVQLVESGGGLVQAGGSLRLSCAAS*GSTLYGYV*MGWYRQAPGKE<br>RELVA*AITDSGGST*YYADSVKGRFTISRDNAKNTVYLQMNSLKP<br>EDTAVYYC*AADYPYFYYVKSYDY*WGQGTQVTVSS |
| B03 | QVQLVESGGGLVQAGGSLRLSCAAS*GRIFSGNA*MGWYRQAPGKE<br>RELVA*AITFSGGST*NYADSVKGRFTISRDNAKNTVYLQMNSLKP<br>EDTAVYYC*NVASSDGYYYFAIGYGY*WGQGTQVTVSS |
| D04’ | QVQLVESGGGLVQAGGSLRLSCAAS*GSTSWRNA*MGWYRQAPGKE<br>RELVA*AISTGGNT*YYADSVKGRFTISRDNAKNTVYLQMNSLKPE<br>DTAVYYC*AAYDASQYGYDY*WGQGTQVTVSS |
| A12’ | QVQLVESGGGLVQAGGSLRLSCAAS*GSTFYYYA*MGWYRQAPGKE<br>RELVA*VISWSGGS*TYADSVKGRFTISRDNAKNTVYLQMNSLKPE<br>DTAVYYC*AAGAYKPVDTSDY*WGQGTQVTVSS |
| C02’ | QVQLVESGGGLVQAGGSLRLSCAAS*GRTFTYNA*MGWYRQAPGKE<br>RELVA*RISFSTGST*YYADSVKGRFTISRDNAKNTVYLQMNSLKP<br>EDTAVYYC*AVSTTSFGAQTGYGPYPYGY*WGQGTQVTVSS |
| D11’ | QVQLVESGGGLVQAGGSLRLSCAAS*GSIFGDNA*MGWYRQAPGKE<br>RELVA*TITFRGAGT*YYADSVKGRFTISRDNAKNTVYLQMNSLKP<br>EDTAVYYC*AKDYWSAWQNNGYDY*WGQGTQVTVSS |
| A05 | QVQLVESGGGLVQAGGSLRLSCAGYAMGWYRQAPGKERELVA*TI*<br>*SGSGGST*YYADSVKGRFTISRDNAKNTVYLQMNSLKPEDTAVYY<br>C*AAAYWDYSYEYYY*WGQGTQVTVSS |

| | |
|---|---|
| C09’ | QVQLVESGGGLVQAGGSLRLSCAAS*GRIFGRNA*MGWYRQAPGKE<br>RELVA*AISWSGGN*TYYADSVKGRFTISRDNAKNTVYLQMNSLKP<br>EDTAVYYC*AKDVSYPYKVTWHYQYDY*WGQGTQVTVSS |
| G01 | QVQLVESGGGLVQAGGSLRLSCAAS*GIIFNGYA*MGWYRQAPGKE<br>RELVA*AITDDGTS*TYYADSVKGRFTISRDNAKNTVYLQMNSLKP<br>EDTAVYYC*AADGGLVDFYY*WGQGTQVTVSS |
| G06 | QVQLVESGGGLVQAGGSLRLSCAAS*GSTFGANT*MGWYRQAPGKE<br>RELVA*AISWSGGT*TYYADSVKGRFTISRDNAKNTVYLQMNSLKP<br>EDTAVYYC*ARYKYPADYGY*WGQGTQVTVSS |
| B11 | QVQLVESGGGLVQAGGSLRLSCAAS*GSTFSSNA*MGWYRQAPGKE<br>RELVA*SINSGDST*YYADSVKGRFTISRDNAKNTVYLQMNSLKPE<br>DTAVYYC*AVVRFGTRPSHIRHTHEYYY*WGQGTQVTVSS |
| B10 | QVQLVESGGGLVQAGGSLRLSCAAS*GSIFVYNA*MGWYRQAPGKE<br>RELVA*AITYSGDDT*YYADSVKGRFTISRDNAKNTVYLQMNSLKP<br>EDTAVYYC*AAEYVEGVLSIYGRSWVYNTYDY*WGQGTQVTVSS |
| A12 | QVQLVESGGGLVQAGGSLRLSCAAS*GRIFSGYA*MGWYRQAPGKE<br>RELVA*TITYTGGST*YYADSVKGRFTISRDNAKNTVYLQMNSLKP<br>EDTAVYYC*NTSPYYVADLRYYY*WGQGTQVTVSS |
| G02 | QVQLVESGGGLVQAGGSLRLSCAAS*GSILLPYA*MGWYRQAPGKE<br>RELVA*TISSSGGST*YYADSVKGRFTISRDNAKNTVYLQMNSLKP<br>EDTAVYYC*AVEQYSNYLENDY*WGQGTQVTVSS |
| B09 | QVQLVESGGGLVQAGGSLRLSCAAS*GSIFVDNA*MGWYRQAPGKE<br>RELVA*SITWRGGRT*SYADSVKGRFTISRDNAKNTVYLQMNSLKP<br>EDTAVYYC*NKVSYRSWYYPAFDY*WGQGTQVTVSS |
| B02’ | QVQLVESGGGLVQAGGSLRLSCAASG*STFTYNT*MGWYRQAPGKE<br>RELVA*SISSTGGST*YYADSVKGRFTISRDNAKNTVYLQMNSLKP<br>EDTAVYYC*AKTGVRARYPYRWGDYDY*WGQGTQVTVSS |
| B02 | QVQLVESGGGLVQAGGSLRLSCAAS*GSTFGLYA*MGWYRQAPGKE<br>RELVA*AITWSGGTT*YYADSVKGRFTISRDNAKNTVYLQMNSLKP<br>EDTAVYYC*AAEVYTYYWYSYYY*WGQGTQVTVSS |
| D08 | QVQLVESGGGLVQAGGSLRLSCAAS*GSTFGTYA*MGWYRQAPGKE<br>RELVA*AISGGGNT*NYADSVKGRFTISRDNAKNTVYLQMNSLKPE<br>DTAVYYCAADWYTYTWGFGYSIYYYWGQGTQVTVSS |
| F10’ | QVQLVESGGGLVQAGGSLRLSCAAS*GSTFWWYA*MGWYRQAPGKE<br>RELVA*TISRGGST*NYADSVKGRFTISRDNAKNTVYLQMNSLKPE<br>DTAVYYC*ARKTRDIRLDY*WGQGTQVTVSS |
| G01’ | QVQLVESGGGLVQAGGSLRLSCAAS*GITFDRYV*MGWYRQAPGKE<br>REFVA*VISRGGRT*YYADSVKGRFTISRDNAKNTVYLQMNSLKPE<br>DTAVYYC*AASVLLYWYWGEDDY*WGQGTQVTVSS |
| B10’ | QVQLVESGGGLVQAGGSLRLSCAAS*GFIFYYNA*MGWYRQAPGKE<br>RELVA*AITWGGGST*YYADSVKGRFTISRDNAKNTVYLQMNSLKP<br>EDTAVYYC*NVLFIAQGSGWTYDY*WGQGTQVTVSS |
| F11’ | QVQLVESGGGLVQAGGSLRLSCAAS*GSTFHHYA*MGWYRQAPGKE<br>RELVA*AITTSGGRT*YYADSVKGRFTISRDNAKNTVYLQMNSLKP<br>EDTAVYYC*AADRFFYRGGYYY*WGQGTQVTVSS |

| | |
|---|---|
| D09 | QVQLVESGGGLVQAGGSLRLSCAAS*GRTFHWNA*MGWYRQAPGKE<br>RELVA*DITSGGST*YYADSVKGRFTISRDNAKNTVYLQMNSLKPE<br>DTAVYYC*AADWFIWWRDDYYAGYDLYDY*WGQGTQVTVSS |
| F02 | QVQLVESGGGLVQAGGSLRLSCAAS*GRIFYSNA*MGWYRQAPGKE<br>RELVA*AITFSGAST*YYADSVKGRFTISRDNAKNTVYLQMNSLKP<br>EDTAVYYC*AAEHYNWVSSYRYYY*WGQGTQVTVSS |
| F01 | QVQLVESGGGLVQAGGSLRLSCAAS*GFIFDSYA*MGWYRQAPGKE<br>RELVA*AITSSGGT*TYYADSVKGRFTISRDNAKNTVYLQMNSLKP<br>EDTAVYYC*AVRSTFRWYGYYY*WGQGTQVTVSS |
| D06 | QVQLVESGGGLVQAGGSLRLSCAAS*GRIFGYYA*MGWYRQAPGKE<br>RELVA*VIRGGVST*NYADSVKGRFTISRDNAKNTVYLQMNSLKPE<br>DTAVYYC*NARRYWAFNAYSKYDY*WGQGTQVTVSS |
| F09' | QVQLVESGGGLVQAGGSLRLSCAAS*GGTFVRYA*MGWYRQAPGKE<br>RELVA*AISSRGDRT*YYADSVKGRFTISRDNAKNTVYLQMNSLKP<br>EDTAVYYC*NTVYYTDSEYDS*WGQGTQVTVSS |
| G04 | QVQLVESGGGLVQAGGSLRLSCAAS*GRIFTGNA*MGWYRQAPGKE<br>REFVA*AISNSGGST*YYADSVKGRFTISRDNAKNTVYLQMNSLKP<br>EDTAVYYC*AASYRVKWKYNY*WGQGTQVTVSS |
| E06' | QVQLVESGGGLVQAGGSLRLSCAAS*GGTIAYNA*MGWYRQAPGKE<br>RELVA*AISSSGGRT*YYADSVKGRFTISRDNAKNTVYLQMNSLKP<br>EDTAVYYC*ATPVTNGFDY*WGQGTQVTVSS |
| H06 | QVQLVESGGGLVQAGGSLRLSCAAS*GSIFYTYA*MGWYRQAPGKE<br>RELVA*AITSTGART*YYADSVKGRFTISRDNAKNTVYLQMNSLKP<br>EDTAVYYC*NALSRAGALKYGGPNDY*WGQGTQVTVSS |
| F10 | QVQLVESGGGLVQAGGSLRLSCAAS*GSTFLLYA*MGWYRQAPGKE<br>RELVA*AISWSGSRT*YYADSVKGRFTISRDNAKNTVYLQMNSLKP<br>EDTAVYYC*AARSGFAGYGY*WGQGTQVTVSS |
| H11 | QVQLVESGGGLVQAGGSLRLSCAAS*GSTFRYYA*MGWYRQAPGKE<br>RELVA*AITRSGAST*YYADSVKGRFTISRDNAKNTVYLQMNSLKP<br>EDTAVYYC*AARSKWNYGRYEY*WGQGTQVTVSS |
| B01 | QVQLVESGGGLVQAGGSLRLSCAAS*GSTFTNYA*MGWYRQAPGKE<br>RELVA*AISNNGGRT*YYADSVKGRFTISRDNAKNTVYLQMNSLKP<br>EDTAVYYC*NARIYQGVYVRWYGY*WGQGTQVTVSS |
| E04' | QVQLVESGGGLVQAGGSLRLSCAAS*GFTFGSYA*MGWYRQAPGKE<br>RELVA*AITISGSST*YYADSVKGRFTISRDNAKNTVYLQMNSLKP<br>EDTAVYYC*NAYWGRGYKTEYYY*WGQGTQVTVSS |
| G02' | QVQLVESGGGLVQAGGSLRLSCAAS*GRIFSRNA*MGWYRQAPGKE<br>RELVA*AITQSGGST*YYADSVKGRFTISRDNAKNTVYLQMNSLKP<br>EDTAVYYC*AAVKYWEYDY*WGQGTQVTVSS |
| H01' | QVQLVESGGGLVQAGGSLRLSCAAS*GRIFGRNT*MGWYRQAPGKE<br>RELVA*AIRSGGST*SYADSVKGRFTISRDNAKNTVYLQMNSLKPE<br>DTAVYYC*NVKSYRYLKDGTFTRKYDY*WGQGTQVTVSS |
| F04' | QVQLVESGGGLVQAGGSLRLSCAAS*GSIFRYYA*MGWYRQAPGKE<br>RELVA*AINSRGTST*YYADSVKGRFTISRDNAKNTVYLQMNSLKP<br>EDTAVYYC*NKLVYRYGSYLEMDY*WGQGTQVTVSS |

| | |
|---|---|
| E10’ | QVQLVESGGGLVQAGGSLRLSCAAS_GRIFSHYA_MGWYRQAPGKE<br>REFVA_AISADGGS_TYYADSVKGRFTISRDNAKNTVYLQMNSLKP<br>EDTAVYYC_AARKYYRTNGY_WGQGTQVTVSS |
| F07’ | QVQLVESGGGLVQAGGSLRLSCAAS_GFTFNRYA_MGWYRQAPGKE<br>RELVA_AISGSGAST_YYADSVKGRFTISRDNAKNTVYLQMNSLKP<br>EDTAVYYC_ATRYSRSYRSRDYYY_WGQGTQVTVSS |
| G09’ | QVQLVESGGGLVQAGGSLRLSCAAS_GITFQRYA_MGWYRQAPGKE<br>RELVA_SISRSGGST_YYADSVKGRFTISRDNAKNTVYLQMNSLKP<br>EDTAVYYC_AARYIVRGGY_WGQGTQVTVSS |
| E05’ | QVQLVESGGGLVQAGGSLRLSCAAS_GYIFVKYA_MGWYRQAPGKE<br>RELVA_AISRSGVRT_YYADSVKGRFTISRDNAKNTVYLQMNSLKP<br>EDTAVYYC_NAYFYANDY_WGQGTQVTVSS |
| F02’ | QVQLVESGGGLVQAGGSLRLSCAAS_GSTFSRNT_MGWYRQAPGKE<br>RELVA_AISKSGGRT_YYADSVKGRFTISRDNAKNTVYLQMNSLKP<br>EDTAVYYC_NAAVYAYASDY_WGQGTQVTVSS |
| D06<br>G26D$_{G27D}$ | QVQLVESGGGLVQAGGSLRLSCAAS_**D**RIFGYYA_MGWYRQAPGKE<br>RELVA_VIRGGVST_NYADSVKGRFTISRDNAKNTVYLQMNSLKPE<br>DTAVYYC_NARRYWAFNAYSKYDY_WGQGTQVTVSS |
| D06<br>T57D$_{T65D}$ | QVQLVESGGGLVQAGGSLRLSCAAS_GRIFGYYA_MGWYRQAPGKE<br>RELVA_VIRGGVS**D**_NYADSVKGRFTISRDNAKNTVYLQMNSLKPE<br>DTAVYYC_NARRYWAFNAYSKYDY_WGQGTQVTVSS |
| D06<br>G26D$_{G27D}$<br>T57D $_{T65D}$ | QVQLVESGGGLVQAGGSLRLSCAAS_**D**RIFGYYA_MGWYRQAPGKE<br>RELVA_VIRGGVS**D**_NYADSVKGRFTISRDNAKNTVYLQMNSLKPE<br>DTAVYYC_NARRYWAFNAYSKYDY_WGQGTQVTVSS |
| D06<br>Y32G$_{Y37G}$ | QVQLVESGGGLVQAGGSLRLSCAAS_GRIFGY**G**A_MGWYRQAPGKE<br>RELVA_VIRGGVST_NYADSVKGRFTISRDNAKNTVYLQMNSLKPE<br>DTAVYYC_NARRYWAFNAYSKYDY_WGQGTQVTVSS |
| D06<br>R52V$_{R57V}$ | QVQLVESGGGLVQAGGSLRLSCAAS_GRIFGYYA_MGWYRQAPGKE<br>RELVA_VI**V**GGVST_NYADSVKGRFTISRDNAKNTVYLQMNSLKPE<br>DTAVYYC_NARRYWAFNAYSKYDY_WGQGTQVTVSS |
| D06<br>Y32G$_{Y37G}$<br>R52V$_{R57V}$ | QVQLVESGGGLVQAGGSLRLSCAAS_GRIFGY**G**A_MGWYRQAPGKE<br>RELVA_VI**V**GGVST_NYADSVKGRFTISRDNAKNTVYLQMNSLKPE<br>DTAVYYC_NARRYWAFNAYSKYDY_WGQGTQVTVSS |
| D06 Y100-<br>$_{Y109-}$ | QVQLVESGGGLVQAGGSLRLSCAAS_GRIFGYYA_MGWYRQAPGKE<br>RELVA_VIRGGVST_NYADSVKGRFTISRDNAKNTVYLQMNSLKPE<br>DTAVYYC_NARR**-**WAFNAYSKYDY_WGQGTQVTVSS |
| D06 T57D<br>$_{T65D}$ Y100-<br>$_{Y109-}$ | QVQLVESGGGLVQAGGSLRLSCAAS_GRIFGYYA_MGWYRQAPGKE<br>RELVA_VIRGGVS**D**_NYADSVKGRFTISRDNAKNTVYLQMNSLKPE<br>DTAVYYC_NARR**-**WAFNAYSKYDY_WGQGTQVTVSS |
| D06<br>R98D$_{R107D}$ | QVQLVESGGGLVQAGGSLRLSCAAS_GRIFGYYA_MGWYRQAPGKE<br>RELVA_VIRGGVST_NYADSVKGRFTISRDNAKNTVYLQMNSLKPE<br>DTAVYYC_NA**D**RYWAFNAYSKYDY_WGQGTQVTVSS |
| D06<br>R99H$_{R108H}$ | QVQLVESGGGLVQAGGSLRLSCAAS_GRIFGYYA_MGWYRQAPGKE<br>RELVA_VIRGGVST_NYADSVKGRFTISRDNAKNTVYLQMNSLKPE<br>DTAVYYC_NAR**H**YWAFNAYSKYDY_WGQGTQVTVSS |

| | |
|---|---|
| D06<br>R98D$_{R107D}$<br>R99H $_{R108H}$ | QVQLVESGGGLVQAGGSLRLSCAASGRIFGYYAMGWYRQAPGKE<br>RELVAVIRGGVSTNYADSVKGRFTISRDNAKNTVYLQMNSLKPE<br>DTAVYYCNARRYWAFNAYSKYDYWGQGTQVTVSS |
| D06<br>A97H$_{A106H}$ | QVQLVESGGGLVQAGGSLRLSCAASGRIFGYYAMGWYRQAPGKE<br>RELVAVIRGGVSTNYADSVKGRFTISRDNAKNTVYLQMNSLKPE<br>DTAVYYCN**H**RRYWAFNAYSKYDYWGQGTQVTVSS |
| D06<br>N96W$_{N106 W}$ | QVQLVESGGGLVQAGGSLRLSCAASGRIFGYYAMGWYRQAPGKE<br>RELVAVIRGGVSTNYADSVKGRFTISRDNAKNTVYLQMNSLKPE<br>DTAVYYC**W**ARRYWAFNAYSKYDYWGQGTQVTVSS |
| D06<br>G54F$_{G59F}$<br>R98D $_{R107D}$ | QVQLVESGGGLVQAGGSLRLSCAASGRIFGYYAMGWYRQAPGKE<br>RELVAVIRG**F**VSTNYADSVKGRFTISRDNAKNTVYLQMNSLKPE<br>DTAVYYCNA**D**RYWAFNAYSKYDYWGQGTQVTVSS |
| D06<br>G26D$_{G27D}$<br>R98D $_{R107D}$ | QVQLVESGGGLVQAGGSLRLSCAAS**D**RIFGYYAMGWYRQAPGKE<br>RELVAVIRGGVSTNYADSVKGRFTISRDNAKNTVYLQMNSLKPE<br>DTAVYYCNA**D**RYWAFNAYSKYDYWGQGTQVTVSS |
| D06<br>G54F$_{G59F}$ | QVQLVESGGGLVQAGGSLRLSCAASGRIFGYYAMGWYRQAPGKE<br>RELVAVIRG**F**VSTNYADSVKGRFTISRDNAKNTVYLQMNSLKPE<br>DTAVYYCNARRYWAFNAYSKYDYWGQGTQVTVSS |
| D06<br>F29R$_{F30R}$ | QVQLVESGGGLVQAGGSLRLSCAASGRI**R**GYYAMGWYRQAPGKE<br>RELVAVIRGGVSTNYADSVKGRFTISRDNAKNTVYLQMNSLKPE<br>DTAVYYCNARRYWAFNAYSKYDYWGQGTQVTVSS |
| D06<br>F29R$_{F30R}$<br>T57D$_{T65D}$ | QVQLVESGGGLVQAGGSLRLSCAASGRI**R**GYYAMGWYRQAPGKE<br>RELVAVIRGGVS**D**NYADSVKGRFTISRDNAKNTVYLQMNSLKPE<br>DTAVYYCNARRYWAFNAYSKYDYWGQGTQVTVSS |
| D06<br>G26D$_{G27D}$<br>A97H $_{A106H}$ | QVQLVESGGGLVQAGGSLRLSCAAS**D**RIFGYYAMGWYRQAPGKE<br>RELVAVIRGGVSTNYADSVKGRFTISRDNAKNTVYLQMNSLKPE<br>DTAVYYCN**H**RRYWAFNAYSKYDYWGQGTQVTVSS |
| D06<br>G26D$_{G27D}$<br>Y32G $_{Y37G}$ | QVQLVESGGGLVQAGGSLRLSCAAS**D**RIFGY**G**AMGWYRQAPGKE<br>RELVAVIRGGVSTNYADSVKGRFTISRDNAKNTVYLQMNSLKPE<br>DTAVYYCNARRYWAFNAYSKYDYWGQGTQVTVSS |
| D06<br>R52V$_{R57V}$<br>T57D $_{T65D}$ | QVQLVESGGGLVQAGGSLRLSCAASGRIFGYYAMGWYRQAPGKE<br>RELVAVI**V**GGVS**D**NYADSVKGRFTISRDNAKNTVYLQMNSLKPE<br>DTAVYYCNARRYWAFNAYSKYDYWGQGTQVTVSS |
| E10’<br>G26D$_{G27D}$ | QVQLVESGGGLVQAGGSLRLSCAAS**D**RIFSHYAMGWYRQAPGKE<br>REFVAAISADGGSTYYADSVKGRFTISRDNAKNTVYLQMNSLKP<br>EDTAVYYCAARKYYRTNGYWGQGTQVTVSS |
| E10’<br>F29R$_{F30R}$ | QVQLVESGGGLVQAGGSLRLSCAASGRI**R**SHYAMGWYRQAPGKE<br>REFVAAISADGGSTYYADSVKGRFTISRDNAKNTVYLQMNSLKP<br>EDTAVYYCAARKYYRTNGYWGQGTQVTVSS |
| E10’<br>G26D$_{G27D}$<br>F29R$_{F30R}$ | QVQLVESGGGLVQAGGSLRLSCAAS**D**RI**R**SHYAMGWYRQAPGKE<br>REFVAAISADGGSTYYADSVKGRFTISRDNAKNTVYLQMNSLKP<br>EDTAVYYCAARKYYRTNGYWGQGTQVTVSS |
| E10’<br>Y32G$_{Y37G}$ | QVQLVESGGGLVQAGGSLRLSCAASGRIFSH**G**AMGWYRQAPGKE<br>REFVAAISADGGSTYYADSVKGRFTISRDNAKNTVYLQMNSLKP<br>EDTAVYYCAARKYYRTNGYWGQGTQVTVSS |

| | |
|---|---|
| E10’<br>T58D$_{T65D}$ | QVQLVESGGGLVQAGGSLRLSCAAS*GRIFSHYA*MGWYRQAPGKE<br>REFVA*AISADGGS***D**YYADSVKGRFTISRDNAKNTVYLQMNSLKP<br>EDTAVYYC*AARKYYRTNGY*WGQGTQVTVSS |
| E10’<br>Y32G$_{Y37G}$<br>T58D$_{T65D}$ | QVQLVESGGGLVQAGGSLRLSCAAS*GRIFSHYA*MGWYRQAPGKE<br>REFVA*AISADGGS***D**YYADSVKGRFTISRDNAKNTVYLQMNSLKP<br>EDTAVYYC*AARKYYRTNGY*WGQGTQVTVSS |
| E10’<br>S52V$_{S57V}$ | QVQLVESGGGLVQAGGSLRLSCAAS*GRIFSHYA*MGWYRQAPGKE<br>REFVA*AI**V**ADGGS*TYYADSVKGRFTISRDNAKNTVYLQMNSLKP<br>EDTAVYYC*AARKYYRTNGY*WGQGTQVTVSS |
| E10’<br>R99D$_{R107D}$ | QVQLVESGGGLVQAGGSLRLSCAAS*GRIFSHYA*MGWYRQAPGKE<br>REFVA*AISADGGS*TYYADSVKGRFTISRDNAKNTVYLQMNSLKP<br>EDTAVYYC*AARKYYRTNGY*WGQGTQVTVSS |
| E10’<br>S52V$_{S57V}$<br>R99D$_{R107D}$ | QVQLVESGGGLVQAGGSLRLSCAAS*GRIFSHYA*MGWYRQAPGKE<br>REFVA*AISADGGS*TYYADSVKGRFTISRDNAKNTVYLQMNSLKP<br>EDTAVYYC*AARKYYRTNGY*WGQGTQVTVSS |
| E10’<br>G55H$_{G62H}$ | QVQLVESGGGLVQAGGSLRLSCAAS*GRIFSHYA*MGWYRQAPGKE<br>REFVA*AISAD**H**GS*TYYADSVKGRFTISRDNAKNTVYLQMNSLKP<br>EDTAVYYC*AARKYYRTNGY*WGQGTQVTVSS |
| E10’<br>G55H$_{G62H}$<br>T58D$_{T65D}$ | QVQLVESGGGLVQAGGSLRLSCAAS*GRIFSHYA*MGWYRQAPGKE<br>REFVA*AISAD**H**GS*TYYADSVKGRFTISRDNAKNTVYLQMNSLKP<br>EDTAVYYC*AARKYYRTNGY*WGQGTQVTVSS |
| E10’<br>Y102E$_{Y110E}$ | QVQLVESGGGLVQAGGSLRLSCAAS*GRIFSHYA*MGWYRQAPGKE<br>REFVA*AISADGGS*TYYADSVKGRFTISRDNAKNTVYLQMNSLKP<br>EDTAVYYC*AARKY**E**RTNGY*WGQGTQVTVSS |
| E10’<br>R99D$_{R107D}$<br>Y102E$_{Y110E}$ | QVQLVESGGGLVQAGGSLRLSCAAS*GRIFSHYA*MGWYRQAPGKE<br>REFVA*AISADGGS*TYYADSVKGRFTISRDNAKNTVYLQMNSLKP<br>EDTAVYYC*AA**D**KY**E**RTNGY*WGQGTQVTVSS |
| E10’<br>A97W$_{A105W}$ | QVQLVESGGGLVQAGGSLRLSCAAS*GRIFSHYA*MGWYRQAPGKE<br>REFVA*AISADGGS*TYYADSVKGRFTISRDNAKNTVYLQMNSLKP<br>EDTAVYYC***W**ARKYYRTNGY*WGQGTQVTVSS |
| E10’<br>G26D$_{G27D}$<br>A97W$_{A105W}$ | QVQLVESGGGLVQAGGSLRLSCAAS***D**RIFSHYA*MGWYRQAPGKE<br>REFVA*AISADGGS*TYYADSVKGRFTISRDNAKNTVYLQMNSLKP<br>EDTAVYYC***W**ARKYYRTNGY*WGQGTQVTVSS |
| E10’<br>A98H$_{A106H}$ | QVQLVESGGGLVQAGGSLRLSCAAS*GRIFSHYA*MGWYRQAPGKE<br>REFVA*AISADGGS*TYYADSVKGRFTISRDNAKNTVYLQMNSLKP<br>EDTAVYYC*A**H**RKYYRTNGY*WGQGTQVTVSS |
| AT118 h32 | EVQLVESGGGLVQPGGSLRLSCAAS*GYIYRRYR*MGWYRQAPGKG<br>REFVA*AISGGSST*NYADSVKGRFTISRDNSKNTVYLQMNSLRAE<br>DTAVYYC*AAYRIVSDPRVY*WGQGTQVTVSS |
| AT118 h32<br>G26D$_{G27D}$ | EVQLVESGGGLVQPGGSLRLSCAAS***D**YIYRRYR*MGWYRQAPGKG<br>REFVA*AISGGSST*NYADSVKGRFTISRDNSKNTVYLQMNSLRAE<br>DTAVYYC*AAYRIVSDPRVY*WGQGTQVTVSS |

| | |
|---|---|
| AT118 h32 Y29R$_{Y30R}$ | EVQLVESGGGLVQPGGSLRLSCAAS*GYI**R**RRYR*MGWYRQAPGKG<br>REFVA*AISGGSST*NYADSVKGRFTISRDNSKNTVYLQMNSLRAE<br>DTAVYYC*AAYRIVSDPRVY*WGQGTQVTVSS |
| AT118 h32 R31D$_{R36D}$ | EVQLVESGGGLVQPGGSLRLSCAAS*GYIYR**D**YR*MGWYRQAPGKG<br>REFVA*AISGGSST*NYADSVKGRFTISRDNSKNTVYLQMNSLRAE<br>DTAVYYC*AAYRIVSDPRVY*WGQGTQVTVSS |
| AT118 h32 Y32G$_{Y37G}$ | EVQLVESGGGLVQPGGSLRLSCAAS*GYIYRR**G**R*MGWYRQAPGKG<br>REFVA*AISGGSST*NYADSVKGRFTISRDNSKNTVYLQMNSLRAE<br>DTAVYYC*AAYRIVSDPRVY*WGQGTQVTVSS |
| AT118 h32 S52V$_{S57V}$ | EVQLVESGGGLVQPGGSLRLSCAAS*GYIYRRYR*MGWYRQAPGKG<br>REFVA*AI**V**GGSST*NYADSVKGRFTISRDNSKNTVYLQMNSLRAE<br>DTAVYYC*AAYRIVSDPRVY*WGQGTQVTVSS |
| AT118 h32 G54F$_{G59F}$ | EVQLVESGGGLVQPGGSLRLSCAAS*GYIYRRYR*MGWYRQAPGKG<br>REFVA*AISG**F**SST*NYADSVKGRFTISRDNSKNTVYLQMNSLRAE<br>DTAVYYC*AAYRIVSDPRVY*WGQGTQVTVSS |
| AT118 h32 T57D$_{T65D}$ | EVQLVESGGGLVQPGGSLRLSCAAS*GYIYRRYR*MGWYRQAPGKG<br>REFVA*AISGGSS**D***NYADSVKGRFTISRDNSKNTVYLQMNSLRAE<br>DTAVYYC*AAYRIVSDPRVY*WGQGTQVTVSS |
| AT118 h32 T57I$_{T65I}$ | EVQLVESGGGLVQPGGSLRLSCAAS*GYIYRRYR*MGWYRQAPGKG<br>REFVA*AISGGSS**I***NYADSVKGRFTISRDNSKNTVYLQMNSLRAE<br>DTAVYYC*AAYRIVSDPRVY*WGQGTQVTVSS |
| AT118 h32 A96W$_{A105W}$ | EVQLVESGGGLVQPGGSLRLSCAAS*GYIYRRYR*MGWYRQAPGKG<br>REFVA*AISGGSST*NYADSVKGRFTISRDNSKNTVYLQMNSLRAE<br>DTAVYYC*__W__AYRIVSDPRVY*WGQGTQVTVSS |
| AT118 h32 A97H$_{A106H}$ | EVQLVESGGGLVQPGGSLRLSCAAS*GYIYRRYR*MGWYRQAPGKG<br>REFVA*AISGGSST*NYADSVKGRFTISRDNSKNTVYLQMNSLRAE<br>DTAVYYC*A**H**YRIVSDPRVY*WGQGTQVTVSS |
| AT118 h32 Y98D$_{Y107D}$ | EVQLVESGGGLVQPGGSLRLSCAAS*GYIYRRYR*MGWYRQAPGKG<br>REFVA*AISGGSST*NYADSVKGRFTISRDNSKNTVYLQMNSLRAE<br>DTAVYYC*AA**D**RIVSDPRVY*WGQGTQVTVSS |
| AT118 h32 R99D$_{R108D}$ | EVQLVESGGGLVQPGGSLRLSCAAS*GYIYRRYR*MGWYRQAPGKG<br>REFVA*AISGGSST*NYADSVKGRFTISRDNSKNTVYLQMNSLRAE<br>DTAVYYC*AAY**D**IVSDPRVY*WGQGTQVTVSS |
| AT118 h32 G26D$_{G27D}$ T57I$_{T65I}$ | EVQLVESGGGLVQPGGSLRLSCAAS*__D__YIYRRYR*MGWYRQAPGKG<br>REFVA*AISGGSS**I***NYADSVKGRFTISRDNSKNTVYLQMNSLRAE<br>DTAVYYC*AAYRIVSDPRVY*WGQGTQVTVSS |
| Cablivi (12A2H1) | EVQLVESGGGLVQPGGSLRLSCAAS*GRTFSYNP*MGWFRQAPGKG<br>RELVA*AISRTGGST*YYPDSVEGRFTISRDNAKRMVYLQMNSLRA<br>EDTAVYYC*AAAGVRAEDGRVRTLPSEYTF*WGQGTQVTVSS |

**Table S2. Orthorep Substitutions**

| Clone | Total # substitutions detected | # substitutions computationally predicted to decrease polyreactivity one-hot | # substitutions computationally predicted to decrease polyreactivity k-mer |
|---|---|---|---|
| E05’ | 240 | 233 | 234 |
| F02’ | 49 | 33 | 32 |
| G09’ | 62 | 43 | 47 |
| F07’ | 14 | 13 | 13 |

### Table S3. Crystallography data collection and refinement statistics

| | AT118i4h32 | AT118i4h32 G26D[27] T57I[65] |
|---|---|---|
| **PDB ID** | 7T83 | 7T84 |
| **Data collection** | | |
| Space group | *P2₁* | *P2₁* |
| Cell dimensions | | |
| *a*, *b*, *c* (Å) | 80.1, 85.4, 84.4 | 78.7, 89.2, 83.1 |
| α, β, γ (°) | 90, 108.6, 90 | 90, 110.1, 90 |
| Resolution (Å) | 31.5 – 2.1 (2.2 – 2.1) * | 46.3 – 1.6 (1.7-1.6) |
| $R_{merge}$ (%) | 6.8 (174.3) | 8.9 (174.9) |
| $CC_{1/2}$ (%) | 99.8 (67.1) | 99.9 (53.6) |
| $I / \sigma I$ | 12.1 (0.84) | 13.4 (1.4) |
| Completeness (%) | 98.9 (97.2) | 98.6 (97.4) |
| Redundancy | 6.8 (6.7) | 6.8 (7.1) |
| **Refinement** | | |
| Resolution (Å) | 31.5 – 2.1 | 46.3 – 1.6 |
| No. reflections | 62627 | 139941 |
| $R_{work}$ / $R_{free}$ (%) | 20.8/25.7 | 16.1 / 18.9 |
| No. atoms | 7428 | 8651 |
| Protein | 7305 | 7691 |
| Ligand | 52 | 139 |
| Water | 71 | 821 |
| *B*-factors | 86.3 | 32.3 |
| Protein | 86.4 | 31.3 |
| Ligand | 107.3 | 41.0 |
| Water | 68.7 | 39.9 |
| R.m.s. deviations | | |
| Bond lengths (Å) | 0.003 | 0.02 |
| Bond angles (°) | 0.9 | 1.62 |

*Values in parentheses are for highest-resolution

**Table S4. Melting temperatures of AT118i4h32 variants**

| | $T_m$ (°C) |
|---|---|
| AT118i4h32 | 64.0 ± 0.05 |
| AT118i4h32 G26D$^{27}$ | 66.3 ± 0.05 |
| AT118i4h32 T57I$^{65}$ | 62.8 ± 0.31 |
| AT118i4h32 G26D$^{27}$ + T57I$^{65}$ | 65.7 ± 0.06 |

Effect of G26D$^{27}$ and T57I$^{65}$ substitutions on melting temperature ($T_m$, °C) of AT118i4h32. Error represents standard error of the mean determined from three independent experiments performed in technical triplicates.

**Table S5. Table of nanobody $K_i$ values**

| | [3H]-Olmesartan / Membrane AT1R |
|---|---|
| Losartan | -7.34 ± 0.07 (45.7 nM) |
| AT118i4h32 | -7.14 ± 0.08 (72.4 nM) |
| AT118i4h32 G26D$^{27}$ | -7.15 ± 0.12 (70.8 nM) |
| AT118i4h32 T57I$^{65}$ | -7.19 ± 0.13 (64.6 nM) |
| AT118i4h32 G26D$^{27}$ + T57I$^{65}$ | -7.10 ± 0.14 (79.4 nM) |

Effects of G26D$^{27}$ and T57I$^{65}$ substitutions on ligand binding. Log Ki values (Ki values shown in parentheses) were determined using the Kd value of [3H]-Olmesartan (Kd 0.29 +/- 0.03 nM). Error represents standard error of the mean from three independent experiments.

**Table S6. Table of nanobody rescue mutation predictions with models trained on deeper FACS sorting experiments**

| Name | Logistic regression onehot CDRS original | Logistic regression 3mer CDRS original | Logistic regression onehot CDRS deep | Logistic regression 3mer CDRS deep | CNN CDRS deep | RNN CDRS deep |
|---|---|---|---|---|---|---|
| AT118 h32 G26D$_{G27D}$ | 1.42 | -1.11 | -0.70 | -0.77 | 0.60 | 0.51 |
| AT118 h32 R31D$_{R36D}$ | 0.94 | 0.33 | 0.09 | 0.71 | 0.69 | 0.27 |
| AT118 h32 Y29R$_{Y30R}$ | 0.96 | -1.29 | -1.15 | -1.41 | 0.52 | 0.55 |
| AT118 h32 Y32G$_{Y37G}$ | 0.81 | -1.33 | 0.40 | -1.19 | 0.47 | 0.55 |
| AT118 h32 G54F$_{G59F}$ | 1.25 | -0.73 | -1.08 | -0.53 | 0.48 | 0.57 |
| AT118 h32 S52V$_{S57V}$ | 1.19 | -0.47 | -0.57 | -0.33 | 0.48 | 0.53 |
| AT118 h32 T57I$_{T65I}$ | 1.20 | -0.71 | -1.04 | -0.27 | 0.40 | 0.54 |
| AT118 h32 A96W$_{A105W}$ | 0.95 | -0.75 | -1.16 | -0.72 | 0.57 | 0.61 |
| AT118 h32 A97H$_{A106H}$ | 0.91 | -0.28 | -0.70 | -0.58 | 0.48 | 0.36 |
| AT118 h32 R99D$_{R108D}$ | 0.78 | -0.69 | -0.15 | -0.07 | 0.61 | 0.73 |
| AT118 h32 Y98D$_{Y107D}$ | 1.03 | 0.22 | -0.33 | -0.11 | 0.61 | 0.65 |
| AT118 h32 | -0.33 | -1.11 | -1.11 | -0.87 | 0.48 | 0.56 |
| E10' F29R$_{F30R}$ | -0.52 | -2.44 | -1.67 | -2.16 | 0.25 | 0.13 |
| E10' G26D$_{G27D}$ | -0.08 | -2.73 | -1.44 | -1.18 | 0.33 | 0.17 |
| E10' G26D$_{G27D}$ F29R$_{F30R}$ | 1.22 | -2.44 | -1.27 | -1.84 | 0.39 | 0.14 |
| E10' G26D$_{G27D}$ A97W$_{A105W}$ | 1.20 | -2.02 | -1.49 | -0.84 | 0.40 | 0.17 |
| E10' Y32G$_{Y37G}$ | -0.69 | -2.40 | -0.33 | -1.02 | 0.23 | 0.17 |

| | | | | | | |
|---|---|---|---|---|---|---|
| E10’ Y32G$_{Y37G}$ T58D$_{T65D}$ | 1.46 | -1.82 | -0.33 | -0.32 | 0.26 | 0.17 |
| E10’ G55H$_{G62H}$ | -0.10 | -2.67 | -2.16 | -1.25 | 0.32 | 0.17 |
| E10’ G55H$_{G62H}$ T58D$_{T65D}$ | 2.04 | -2.09 | -2.16 | -0.56 | 0.31 | 0.18 |
| E10’ S52V$_{S57V}$ | -0.31 | -1.67 | -1.30 | -1.23 | 0.28 | 0.15 |
| E10’ S52V$_{S57V}$ R99D$_{R107D}$ | 1.25 | 0.22 | -0.21 | 0.06 | 0.67 | 0.17 |
| E10’ T58D$_{T65D}$ | 0.33 | -2.15 | -1.84 | -0.80 | 0.23 | 0.16 |
| E10’ A97W$_{A105W}$ | -0.54 | -2.02 | -1.89 | -1.15 | 0.29 | 0.18 |
| E10’ A98H$_{A106H}$ | -0.59 | -1.84 | -1.44 | -0.90 | 0.25 | 0.18 |
| E10’ R99D$_{R107D}$ | -0.27 | -0.84 | -0.74 | -0.20 | 0.55 | 0.17 |
| E10’ R99D$_{R107D}$ Y102E$_{Y110E}$ | 0.80 | -0.07 | -0.12 | 1.04 | 0.73 | 0.24 |
| E10’ Y102E$_{Y110E}$ | -0.75 | -1.96 | -1.21 | -0.26 | 0.38 | 0.17 |
| E10’ | -1.82 | -2.73 | -1.84 | -1.50 | 0.21 | 0.15 |
| D06 F29R$_{F30R}$ | 1.49 | -1.34 | -0.71 | -1.87 | 0.27 | 0.18 |
| D06 F29R$_{F30R}$ T57D$_{T65D}$ | 3.64 | -0.85 | -0.71 | -1.70 | 0.43 | 0.27 |
| D06 G26D$_{G27D}$ | 1.93 | -1.07 | -0.47 | -0.93 | 0.35 | 0.24 |
| D06 G26D$_{G27D}$ Y32G $_{Y37G}$ | 3.07 | -0.61 | 1.04 | -0.60 | 0.37 | 0.20 |
| D06 G26D$_{G27D}$ T57D $_{T65D}$ | 4.08 | -0.58 | -0.48 | -0.76 | 0.41 | 0.16 |
| D06 G26D$_{G27D}$ A97H $_{A106H}$ | 3.17 | -0.46 | -0.07 | -1.00 | 0.38 | 0.24 |
| D06 G26D$_{G27D}$ R98D $_{R107D}$ | 3.49 | 0.68 | 0.62 | 0.15 | 0.61 | 0.36 |
| D06 Y32G$_{Y37G}$ | 1.32 | -0.61 | 0.63 | -0.88 | 0.29 | 0.19 |

| | | | | | | |
|---|---|---|---|---|---|---|
| D06 Y32G$_{Y37G}$ R52V$_{R57V}$ | 2.83 | 0.00 | 1.47 | -0.05 | 0.65 | 0.30 |
| D06 G54F$_{G59F}$ | 1.77 | -0.55 | -0.85 | -0.69 | 0.27 | 0.24 |
| D06 G54F$_{G59F}$ R98D $_{R107D}$ | 3.32 | 1.19 | 0.24 | 0.39 | 0.71 | 0.30 |
| D06 R52V$_{R57V}$ | 1.70 | -0.46 | -0.04 | -0.38 | 0.59 | 0.28 |
| D06 R52V$_{R57V}$ T57D $_{T65D}$ | 3.84 | 0.03 | -0.04 | -0.21 | 0.66 | 0.19 |
| D06 T57D$_{T65D}$ | 2.34 | -0.58 | -0.88 | -1.04 | 0.40 | 0.24 |
| D06 T57D $_{T65D}$ Y100- $_{Y109-}$ | 3.13 | 0.14 | -0.54 | -1.14 | 0.46 | 0.25 |
| D06 A97H$_{A106H}$ | 1.42 | -0.46 | -0.48 | -1.28 | 0.28 | 0.24 |
| D06 N96W$_{N106W}$ | 1.36 | -0.85 | -1.02 | -1.17 | 0.28 | 0.24 |
| D06 R98D$_{R107D}$ | 1.74 | 0.68 | 0.22 | -0.13 | 0.70 | 0.33 |
| D06 R99H$_{R108H}$ | 2.81 | 1.38 | 0.80 | 0.33 | 0.78 | 0.61 |
| D06 R98D$_{R107D}$ R99H $_{R108H}$ | 1.26 | 0.14 | -0.29 | -0.68 | 0.34 | 0.32 |
| D06 Y100- $_{Y109-}$ | 0.98 | -0.37 | -0.54 | -1.32 | 0.29 | 0.30 |
| D06 | 0.19 | -1.07 | -0.88 | -1.21 | 0.26 | 0.26 |

### ONLINE METHODS

#### Generation of insect cell membrane polyreactivity reagent

Insect cell membrane polyreactivity reagent was generated as described previously [1]. Briefly, 250 mL of sf9 insect cells at a density of $4x10^6$ cells/mL were pelleted, washed in 100 mL PBS + 1% BSA followed by 30 mL Buffer B (50 mM Hepes pH 7.2, 150 mM NaCl, 2 mM $CaCl_2$, 5 mM KCl, 5 mM $MgCl_2$, 10% glycerol). The cell pellet was resuspended in 3x pellet volume of Buffer B with a protease inhibitor tablet (Roche) and lysed with a dounce homogenizer. The membrane fraction was pelleted by centrifugation at 40,000xg for 1 hour, washed with 1 mL Buffer B and resuspended in 3 mL of Buffer B with dounce homogenization. Total protein was quantified using the DC protein assay (Biorad) following manufactures instructions. The membrane fraction was diluted to ~1 mg/mL in Buffer B and biotinylated with 200 µM NHS-LC-Biotin for three hours at 4°C. 20 mM Tris pH 8 was added to quench excess NHS-LC-Biotin. The biotinylated membrane fraction was centrifuged at 40,000xg for 1 hour and the pellet was washed 5 times with Buffer B and resuspended in 3 mL Buffer B + 10% glycerol by dounce homogenization, and total protein was quantified by the DC protein assay (Biorad). The membrane fraction was diluted to 1 mg/mL in solubilization buffer (50 mM Hepes pH 7.2, 150 mM NaCl, 2 mM $CaCl_2$, 5 mM KCl, 5 mM $MgCl_2$, 10% glycerol, 1% DDM, 1x protease inhibitor pH 7.2) and stirred overnight at 4°C. The mixture was centrifuged for 40,000xg for 1 hour. Total protein in the supernatant containing the solubilized membrane fraction was quantified using the DC protein assay and aliquots were flash frozen and stored at -80°C.

#### Yeast Sorting

A yeast surface display library containing >2x$10^9$ synthetic nanobody sequences where each amino acid position is diversified based on the natural llama immunological repertoire [2]. Nanobodies are tethered to the yeast cell surface on a synthetic stalk [3] from a vector encoding nourseothricin resistance [2] in *Saccharomyces cerevisiae* BJ5465. Nanobody expression was induced for 36-48 hours in dropout medium without tryptophan (-Trp) supplemented with galactose. 5x$10^9$ yeast cells were stained with 10% PSR in selection buffer (20 mM Hepes pH 7.5, 100 mM NaCl, 0.1% DDM, 0.01% CHS 0.05% BSA, 5 mM CaCl2, 10 mM maltose) at 4°C for 1 hour. Cells were spun down, resuspended in 4.5 mL selection buffer, and incubated with 500 µL streptavidin conjugated microbeads (Miltenyi) for 20 min at 4°C. Cells were washed with 5 mL selection buffer and applied to a LS column (Miltenyi). The column was washed with 8 mL selection buffer. 2.8x$10^7$ yeast clones were collected in the MACS elution and subjected to a round of FACS. 5x$10^7$ yeast cells were stained with 10% PSR in selection buffer for 1 hour at 4°C, washed with selection buffer, and stained with a 1:100 dilution of Alexaflour-647 conjugated anti-HA antibody to detect nanobody expression and Alexaflour-488 conjugated streptavidin (Biolegend) to detect biotinylated PSR positive cells for 15 min. Cells were washed with selection buffer and resuspended in selection buffer for FACS on a SONY SH800 cell sorter. Gates to detect low and high polyreactivity clones were set based upon cells stained with Alexaflour-647 conjugated anti-HA antibody and Alexaflour-488 conjugated streptavidin. For initial experiments 5x$10^7$ total yeast cells were sorted. 3.7x$10^6$ low polyreactivity clones and the most polyreactive clones (top ~1% containing 3.2 x$10^6$ clones) were collected. To obtain additional sequencing data the MACS enriched library was subjected to additional rounds of cell sorting to collect 4.6x$10^7$ highly polyreactive clones and 9.8 x$10^5$ low polyreactivity clones. Cells from the MACS elution and low and high poyreactivity FACS sorted populations were plated on -Trp media to obtain single clones. Flow cytometry gating figures were generated in FlowJo (10.8.1).

**Deep sequencing**

The nanobody sequences were amplified from the low and high polyreactivity populations via colony PCR. Media was aspirated from 4x$10^6$ pelleted yeast cells. Cells were microwaved for 1 min on high power twice. Cells were resuspended in 1x Q5 High-Fidelity master mix containing 0.3 µM forward (GTTCAATTGGACAAGAGAGAAGCT) and reverse primers (GTAATCTGGAACATCGTATGGGTA). Cells were subjected to a 4 min incubation at 95°C and DNA was amplified following the manufacturers protocol. Amplified DNA was gel extracted and evaluated via Illumina MiSeq in a 2 x 250 paired-end sequencing reaction.

**NGS analysis and sequence processing**

Fastq sequences from deep sequencing were processed using the FastQC, Trimmomatic, FASTX-Toolkit programs. Sequences were translated to protein sequences using the Biopython package and only nanobody sequences were retrieved by selecting for the highly conserved final beta strand sequence. The nanobody sequences were aligned using ANARCI with standard IMGT numbering to identify the CDR regions. For our dataset of sequences to train the supervised models, we limited nanobody sequences to sequences with a CDR1 length of 8, a CDR2 length of 8 or 9 (9 or 10 in the deeper sequencing exploration, when we include an additional position at the end of CDR2 to include more variability), and CDR3 lengths between 6 and 22. These processing steps leave us with 65,147 unique low polyreactivity sequences and 69,155 unique highly polyreactive sequences that contained 51,308 and 59,623 distinct CDR regions.

**Supervised model development**

The CDR regions were used to build four different types of supervised models: a one-hot logistic regression model, a k-mer logistic regression model, a CNN, and an RNN. The logistic regression models were built using the scikit-learn python package. For the one-hot logistic regression model and CNN model, the sequences were processed into aligned one-hot encoding vectors of amino acids per position (via IMGT numbering). For the RNN, sequences were processed into non-aligned one-hot encoding vectors (padded at the ends of sequences to the longest length). For the k-mer logistic regression model, sequences were processed into vectors of k-mers ranging from single amino acids ("1-mer") to 3-mer motifs. The CNN and RNN models were written in pytorch. The CNN has three convolutional layers (first layer: 1D-convolution (channel dimension size 20 → 32) with kernel size of 3, BatchNorm, and ReLU; second layer: 1D-convolution (channel dimension size 32 → 64) with kernel size of 3, BatchNorm, ReLU, and MaxPool with kernel size of 3, stride of 3; 1D-convolution (channel dimension size 64 → 128) with kernel size of 3, BatchNorm, ReLU, and MaxPool with kernel size of 3, stride of 1) followed by a fully connected layer and a final sigmoid for binary classification and was trained using the Adam optimizer. The RNN has two layers and a hidden size of 128. For splitting sequences by clusters and sequence identity sci-kit learn KMeans clustering and python-levenshtein package was used.

**Yeast plasmid transformation**

pYDS plasmids were transformed into yeast following standard protocol [4]. 2x YPAD media was inoculated with *Saccharomyces cerevisiae* BJ5465 and grown to a density of $2x10^7$ cells / mL. Cells were harvested by centrifugation and washed with water five times. $1E^6$ cells were suspended in a 360 uL transfection mix containing 33.3% PEG3350, 100 mM lithium acetate, 0.28 mg/mL salmon sperm carrier DNA, and 1 μg of the pYDS plasmid encoding the nanobody, synthetic stalk, and nourseothricin resistance cassette. The transformation mixture was incubated at 42°C for 40 minutes. Yeast cells were isolated by centrifugation, washed with water, and resuspended in YPAD. After a 1-2 hour outgrowth at 30°C without shaking, cells were plated on YPAD supplemented with 100 μg/mL nourseothricin.

**Anti-$V_{HH}$ Antibody Staining**

Polyreactivity index panel yeast were grown in -Trp + Glu media for two days at 30°C and induced in -Trp + Gal media at 25°C for two days. After induction, $1x10^6$ yeast cells were washed with DDM selection buffer and were stained with a 1:100 dilution of Alexafluor-488 conjugated Monorab Rabbit Anti-Camelid $V_{HH}$ Antibody (Genescript) and 1:100 dilution of Alexafluor-647 conjugated anti-HA antibody. Following an additional wash, analytical staining was performed using a BD Accuri C6 flow cytometer.

**Recombinant Nanobody Expression and Purification**

Recombinant nanobodies containing a C-terminal V5 epitope and hexahistidine tag were cloned into pET26b and amino acid substitutions were introduced using the QuikChange lighting site-directed mutagenesis kit (Agilent). Plasmids were transformed into *E. coli* BL21(DE3) in Terrific Broth (RPI) supplemented with 4% glycerol and 50 ug/mL kanamycin to an OD600 of 1-2 at 37°C and cooled to 17-20°C for one hour. Protein expression was induced using 0.2 mM isopropyl β-D-1-thiogalactopyranoside (IPTG, Gold Biotechnology) overnight. Bacterial pellets were resuspended in room temperature SET lysis buffer (200 mM Tris pH 8, 500 mM sucrose, 500 μM EDTA) with gentle stirring for 20 minutes, followed by addition of 2x volume ice cold DI $H_2O$, 5 mM magnesium chloride, and 1 μL benzonase nuclease (Sigma-Aldrich) for 1 hour. Cellular debris was removed by centrifugation at 14,000xg for 30 minutes. Following centrifugation, 100 mM sodium chloride was added to the supernatant with stirring for 15 minutes and the supernatant was filtered using glass microfiber filters (Fisher Scientific). Clarified lysate was passed over Protein A resin (Gold Biotechnology) equilibrated with Protein A wash buffer (10 mM sodium phosphate pH 7.5, 100 mM sodium chloride). Then, the column was washed with 10 column volumes Protein A wash buffer, and nanobody was eluted using 10 columns Protein A elution buffer (100 mM sodium phosphate pH 2.5, 100 mM sodium chloride) directly into 1 column volume 2M Hepes pH 8. The Protein A column eluate was passed over a Ni-NTA (Qiagen) column equilibrated with Ni-column wash buffer (20 mM Hepes pH 7.5, 150 mM sodium chloride). The column was then washed with 10 column volumes Ni-NTA wash buffer and eluted using 10 column volumes Ni-NTA elution buffer (20 mM Hepes pH 7.4, 150 mM sodium chloride, 200-400 mM imidazole). The eluate was then dialyzed overnight against SEC buffer (20 mM Hepes pH 7.5, 150 mM sodium chloride, 10% glycerol) and concentrated. Index set nanobodies were purified by size exclusion chromatography using a Superdex S-75 10/300 GL column (GE Healthcare) gel filtration system. Protein purity was assessed by SDS-PAGE.

**Nanobody Polyreactivity ELISA Assays**

Direct ELISA assays were performed similarly to those reported previously. Briefly, high-binding Costar 96-well plates (Corning) were coated with 0.5 μg salmon sperm ssDNA (Abcam), calf thymus dsDNA (Sigma-Aldrich), lipopolysaccharide from *E. coli* (Sigma-Aldrich), chicken egg white lysozyme (Sigma-Aldrich), or 0.25 μg insulin (Fitzgerald) and incubated overnight at 4°C. The next morning, plates were washed three times using wash buffer (PBS pH 7.5, 0.001% Tween), were blocked using blocking buffer (PBS pH 7.5, 0.1% Tween-20, 1 mM EDTA, 2% BSA) for two hours at room temperature, and then were washed three times with wash buffer. Following blocking, nanobodies (200 μL) were incubated at the indicated concentrations at room temperature in PBS pH 7.5 for two hours. After three more washes, plates were incubated with HRP-anti V5 antibody (Abcam ab1325, 1:10,000 dilution) in PBS + 2% BSA for one hour at room temperature. Plates then were washed three times with wash buffer and 1-Step ABTS substrate solution (100 μL, Thermo Scientific) was added to the plates, which were then incubated in the dark for 20 minutes. Stop solution (1% SDS in PBS, 100 μL) was added to each plate and absorbance at 405 nm was measured using a Spectromax M5 microplate reader. Results were analyzed in GraphPad Prism.

**AHEAD Orthogonal Replication**

Nanobodies were amplified using primers PSR_Nb_F and PSR_Nb-R and cloned into the AHEAD integration plasmid (pAW240). The plasmids were linearized with Sca1 and transformed into the AHEAD base strain as previously described[5]. At each cycle, $5x10^7$ cells were labeled with biotinylated insect cell membrane polyreactivity reagent and an HA epitope tag binding antibody as described above and subjected to FACS selection applying a gate that enriches for cells with reduced binding to PSR. The typical number of cells that were selected at each round was 400 out of $2x10^7$ sorted cells. The selected cells were sorted into 3 mL of SC – HLUW media and grown at 30°C with 250 RPM shaking for 48 hours until saturation. Cells cultures were then induced for nanobody display by diluting them at a 1:20 ratio into SC -HLUW media containing 2% galactose instead of glucose and incubated at 20°C for 48 hours. In preparation of next-generation sequencing, p1 plasmid was extracted, as previously described[6], from yeast cultures after the FACS step of each AHEAD cycle. PCRs were performed with Q5 Master Mix (New England Biolabs Cat# M0492S) and primers NGS_p1_F and NGS_p1_R. Following PCR reactions, samples were PCR purified. Amplicon sequencing was performed by the Genewiz and the resulting sequences were analyzed using the methods described above.

**Polyspecifity Reagent Analytical Staining**

Mutations in D06, E10', and AT118i4h32 were introduced using the Quikchange Lightning mutagenesis kit (Agilent), and yeast were transformed using a standard transformation protocol. Polyreactive nanobody panel and mutant yeast were grown in -Trp + Glu media for two days at 30°C and induced in -Trp + Gal media at 25°C for two days. $1x10^6$ yeast were washed with DDM selection buffer, and were stained with a 1:10 dilution of either insect cell PSR reagent or Expi cell PSR reagent for 30 minutes at 4°C with shaking. Following incubation with PSR reagent, yeast were washed with DDM selection buffer and were stained with a 1:100 dilution of Alexafluor-647 conjugated anti-HA antibody and 1:100 dilution of Alexafluor-488 conjugated streptavidin (Biolegend) for 15 minutes at 4°C with shaking. Cells were washed once more with DDM selection buffer and analytical staining was performed using a BD Accuri C6 flow cytometer.

**AT1R Binding Assay**

Expi293F cells stably expressing the tetracycline repressor[7] were stably transfected with a wild-type human FLAG-AT1R containing plasmid (pCDNA Zeo-TetO) to create an inducible cell line, as previously described[8]. Expi293F TetR Zeo FLAG-AT1R cells were grown to 1.5-2x$10^6$ cells/mL induced with 0.4 µg/mL doxycycline hyclate for 24 hours. Cells were washed with cold flow assay buffer (HBS + 0.1% BSA). 2.8x$10^5$ cells were plated and stained in 20 nM of each AT118i4h32 variant in HBS + 0.1% BSA 100 µL reaction volumes for 1 hour at 4ºC with gentle shaking. Cells were washed 2 times with flow assay buffer and subsequently stained with 100 nM of Alexaflour 488 conjugated M1-anti FLAG antibody and 1:200 of Alexaflour 647 conjugated anti-V5 antibody (ThermoFisher) in flow assay buffer + 1 mM $CaCl_2$ (100 µL reaction volumes) for 20 minutes at 4ºC. Cells were washed once and resuspended in flow assay buffer + 1 mM $CaCl_2$. Samples were analyzed on an BD Accuri C6 flow cytometer. Cells were gated for M1-positive singlets. Data were analyzed with BD Accuri C6 Plus software.

**AT1R Signaling Assay**

Expi293F cells stably expressing a tetracycline inducible wild-type human FLAG-AT1R were diluted to 1.5-2 x $10^6$ cells/mL and induced with 0.4 μg/mL doxycycline hyclate for 24-28 hours. 2 x $10^4$ cells were plated into a low-volume 96-well plate, treated with 5 μM of each AT118i4h32 variant for 30 min at 37°C, and stimulated with AngII for 1 hr at 37°C. IP1 was detected with the IP-One Gq kit (CisBio) and read on a SpectraMax M5e plate reader (Molecular Devices).

**Radioligand Binding Assays**

Cell membranes for radioligand binding experiments were prepared from Expi293F cells stably expressing tetracycline inducible wild-type human FLAG-AT1R. AT1R expression was induced at 2x$10^6$ cells/mL with 0.4 μg/mL doxycycline hyclate for 30 hours. Cells were pelleted and washed with cold HBS. Cells were resuspended in 2.5 mL of 20 mM Tris pH 7.4 per gram of cell pellet with a protease inhibitor tablet and lysed by dounce homogenization (100x). Membranes were isolated by centrifugation at 50,000 x g for 20 min. Membranes were resuspended in 2.5 mL of 50 mM Tris pH 7.4, 12.5 mM $MgCl_2$, 150 mM NaCl, 0.2% BSA + protease inhibitor table by dounce homogenization, flash frozen in liquid $N_2$, and stored at -80ºC.

Membranes were incubated with nanobodies and 2 nM [$^3$H]-olmesartan (American Radiolabeled Chemicals) in 50 mM Tris pH 7.4, 12.5 mM MgCl2, 150 mM NaCl, 0.2% BSA for 90 minutes at room temperature. Reactions were harvested on a GF/B filter soaked in water on a 96-well Brandel harvester and washed three times with cold water. Radioligand affinity was measured by saturation binding of [$^3$H]-olmesartan in the presence and absence of 10 μM candesartan. Inhibitory constant ($K_i$) values were determined through a one-site competition binding model in GraphPad Prism. Data represents the mean and SE of three independent biological replicates performed in triplicate.

**Protein Crystallization and Structure Determination**

AT118i4h32 with a N-terminal methionine and alanine and C-terminal His-tag was crystallized at 20°C by sitting drop vapor diffusion from a 1:0.5 µL mixture of protein stock (10 mg/mL AT118i4h32 in 20 mM Hepes pH 7.4, 100 mM NaCl) and reservoir solution (16% PEG 4000, 10% isopropanol, 0.1 M sodium citrate pH 5.6). Crystals were flash cooled directly from the drop in liquid $N_2$.

Diffraction data were collected at 100K on GM/CA beamline 23ID-D at the Advanced Photon Source (APS) at Argonne National Laboratory. Diffraction data were processed using XDS [9]. A camelid antibody 1YC7 was used to solve the structure of AT118i4h32 by molecular replacement using Phaser in the Phenix software suite[10]. The model was rebuilt using Autobuild and manually completed by iterative rounds of model building and refinement using Coot and Phenix.refine with 56 translation/liberation/screw groups. The structure was validated using Molprobity.

AT118i4h32 $G26D^{27}$, $T57I^{65}$ with a N-terminal methionine and alanine and C-terminal His-tag was crystallized at 20°C by sitting drop vapor diffusion from a 0.5:1 µL mixture of protein stock (6.96 mg/mL AT118i4h32 $G26D^{27}$, $T57I^{65}$ in 20 mM Hepes pH 7.4, 150 mM NaCl) and reservoir solution (30% PEG 3350, 280 mM lithium citrate tribasic). Crystals were cryoprotected in 25% PEG 3350, 280 mM lithium citrate tribasic, 15% glycerol and then flash cooled in liquid $N_2$.

Diffraction data were collected at 100K on GM/CA beamline 23ID-D at the Advanced Photon Source (APS) at Argonne National Laboratory. Diffraction data were processed using XDS[9]. The structure of AT118i4h32 was used to solve the AT118i4h32 G26D[27], T57I[65] crystal structure by molecular replacement using Phaser in the Phenix software suite[10]. The model was manually completed by iterative rounds of model building and refinement using Coot and Phenix.refine with 69 translation/liberation/screw groups. The structure was validated using Molprobity. Figures were prepared in PyMol[11]. All software was accessed through SBGrid[12].

**Thermal Shift Assay**

Differential scanning fluorimetry (DSF) experiments were carried out using a Quant Studio 6 real-time PCR machine (Applied Biosystems). 0.1 mg/mL of AT118i4h32 variants in HBS + 10% glycerol was mixed with Protein Thermal Shift Dye (Applied Biosystem) in a 1:100 (v/v) ratio of protein to dye. Samples were heated from 25-90 °C at a rate of 3 °C per minute. The fluorescence was detected with 470 +/- 15 nm excitation and 586 +/- 10 nm emission filters. All samples were measured three biological replicates of technical triplicates. Fluorescence values were fit to the Boltzman equation and melting temperatures (Tm) were extracted from the inflection points of the curves in the Protein Thermal Shift Software (Applied Biosystems).
